# The strand exchange domain of tumor suppressor PALB2 is intrinsically disordered and promotes oligomerization-dependent DNA compaction

**DOI:** 10.1101/2023.06.01.543259

**Authors:** Yevhenii Kyriukha, Maxwell B. Watkins, Jennifer M. Redington, Reza Dastvan, Vladimir N. Uversky, Jesse B. Hopkins, Nicola Pozzi, Sergey Korolev

**Affiliations:** Edward A. Doisy Department of Biochemistry and Molecular Biology, Saint Louis University School of Medicine, St Louis, MO; The Biophysics Collaborative Access Team (BioCat), Departments of Biology and Physics, Illinois Institute of Technology, Chicago, IL; Department of Molecular Medicine and USF Health Byrd Alzheimer’s Research Institute, Morsani College of Medicine, University of South Florida, Tampa, FL

## Abstract

The Partner and Localizer of BRCA2 (PALB2) is a scaffold protein that links BRCA1 with BRCA2 to initiate homologous recombination (HR). PALB2 interaction with DNA strongly enhances HR efficiency in cells. The PALB2 DNA-binding domain (PALB2-DBD) supports strand exchange, a complex multistep reaction conducted by only a few proteins such as RecA-like recombinases and Rad52. Using bioinformatics analysis, small-angle X-ray scattering, circular dichroism, and electron paramagnetic spectroscopy, we determined that PALB2-DBD is an intrinsically disordered region (IDR) forming compact molten globule-like dimer. IDRs contribute to oligomerization synergistically with the coiled-coil interaction. Using confocal single-molecule FRET we demonstrated that PALB2-DBD compacts single-stranded DNA even in the absence of DNA secondary structures. The compaction is bimodal, oligomerization-dependent, and is driven by IDRs, suggesting a novel strand exchange mechanism. Intrinsically disordered proteins (IDPs) are prevalent in the human proteome. Novel DNA binding properties of PALB2-DBD and the complexity of strand exchange mechanism significantly expands the functional repertoire of IDPs. Multivalent interactions and bioinformatics analysis suggest that PALB2 function is likely to depend on formation of protein-nucleic acids condensates. Similar intrinsically disordered DBDs may use chaperone-like mechanism to aid formation and resolution of DNA and RNA multichain intermediates during DNA replication, repair and recombination.

## Introduction

Homologous recombination (HR) is essential for the exchange of genetic information between maternal and paternal alleles and proper chromosome segregation during meiosis, for non-mutagenic repair of large chromosome aberrations such as double-stranded (ds) DNA breaks (DSB), for repair of stalled replication forks, and for a plethora of other DNA transactions (1–4). HR is almost exclusively supported by a single family of RecA-like recombinases, including phage UvsX, bacterial RecA, archaea RadA, and eukaryotic Rad51 and DMC1 (5). The sequences and structures of these RecA-like recombinases are highly conserved, owing to the complexity of a multistep HR reaction involving the formation of an ATP-dependent protein filament on single-stranded (ss) DNA that is capable of searching for a homologous dsDNA and catalyzing strand displacement and strand exchange (3, 6–8). The complexity of this mechanism limits the evolution of more specialized recombinases. However, numerous partner proteins have evolved to regulate organism- and pathway-specific activities of RecA-like recombinases at multiple levels (9–13). RecA-like recombinases recombine long DNA regions with high fidelity but are not efficient for strand exchange between short DNA fragments or between DNA and RNA which may be required during replication fork reversal or transcription/replication collision. Several other protein families are reported to support DNA-pairing and strand exchange, including exonuclease complexes with DNA-annealing proteins such as *E. coli* RecE/RecT, bacteriophage λ Redα/Redβ, and viral UL12/ICP8 complexes(8, 14–16). Another protein capable of strand exchange is the eukaryotic Rad52 (17–19), which is also a major positive regulator of Rad51 in yeast(20, 21). The described proteins form ring-shaped oligomers (22, 23). Rad52 forms an oligomeric toroidal ring that binds ssDNA and dsDNA to promote promiscuous strand exchange (24–26). Rad52 is more efficient than Rad51 in strand exchange between RNA and DNA and is proposed to repair breaks that occur during collision of replication with transcription using newly transcribed mRNA as a homologous template (17, 27). The necessity to process complex multichain DNA and RNA intermediates that are formed in chromatin during all major nucleic acid metabolism reactions is further underscored by limited reports of other proteins with strand annealing and exchange activities. These families include the FANCA protein of the Fanconi anemia (FA) complex, which is critical for DNA interstrand crosslink repair (28), and the RNA-processing FET proteins such as pro-oncoprotein TLS/FUS and the human splicing factor PSF/hPOMp100 (29–31). We discovered that the Partner And Localizer of BRCA2 (PALB2) protein facilitates DNA and RNA strand exchange (32). Human PALB2 is a scaffold protein (1186 amino acids) that links BRCA1 and BRCA2 during HR initiation, where BRCA2 stimulates RAD51 filament formation (33–39). PALB2 interacts with numerous other chromatin and transcription factors, and with RAD51 and its paralogs (34). The N-terminal domain encompassing amino acids 1–195 interacts with DNA (referred to hereafter as PALB2-DBD) (**Fig. 1A**) (40, 41). Alanine substitution of major DNA-binding amino acids 146-RRKK-149 decreases radiation-induced RAD51 foci and HR in cells by 50% (32). We discovered that PALB2-DBD alone promotes RAD51-independent strand exchange *in vitro* (32). PALB2 lacks any sequence or similarities with Rad51 or Rad52, and likely uses a novel mechanism for strand exchange.

**Fig. 1.**
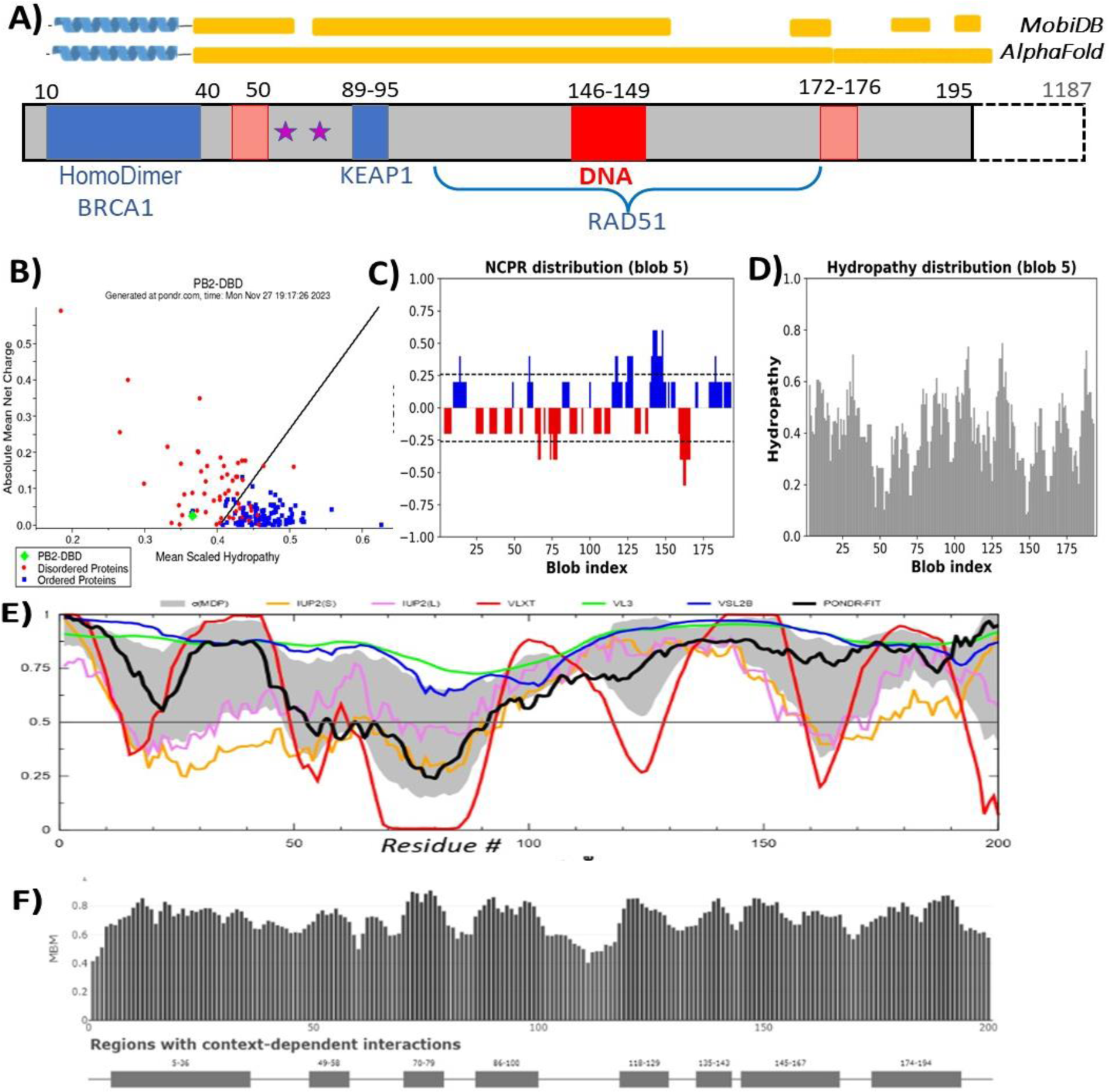
Amino acid sequence features of PALB2-DBD. **(A)** PALB2 sequence motifs. Amino acid sequence numbers are shown above each domain. Shadowed area, PALB2-DBD; blue area, known protein interaction sites; red area, major DNA-binding site; pink area, minor DNA-binding sites; stars, phosphorylation sites; yellow bars above, disordered regions according to MobiDB or AlphaFold prediction. **(B)** Charge-hydropathy plot generated using PONDR® server (www.pondr.com). **(C,D)** CIDER analysis (http://157.245.85.131:8000/CIDER/) of charge distribution (C) and hydrophobicity (D). **(E)** Plot of disorder score against residues number using RIDAO server accordingly to following predictors: mean disorder profile (MDP) error; PONDRs VLXT, VL3 and VSL2, IUPpred-Long, IUPred-Short and PONDR-FIT. **F)** Multiplicity of binding modes evaluated by FuzDrop (https://fuzdrop.bio.unipd.it/predictor).

Here, we present evidence that PALB2-DBD is an intrinsically disordered domain (IDD) or region (IDR) forming a compact dimer. IDRs contribute to dimerization synergistically with the coiled-coil interaction and promote DNA-dependent formation of higher oligomers. Single-stranded (ss) DNA binding further compacts the structure and ssDNA itself is condensed upon binding even in the absence of secondary structure elements. The described oligomerization-dependent DNA condensation and previously reported strand exchange activity by the same domain significantly expands the structural and functional repertoire of intrinsically disordered proteins (IDPs), which represent a third of the human proteome (42). Approximately 30% and 50% of protein and RNA chaperones are estimated to form IDRs, and the proposed strand exchange mechanism will have broad implications as only a few have been mechanistically characterized (43–45). Many eukaryotic DNA repair proteins such as BRCA1 and BRCA2 contain DNA-binding IDRs of unknown function. We hypothesize that such regions represent a novel functional class of DNA repair effectors that can aid transactions between DNA and RNA strands to form or resolve local transient multistrand intermediates that occur near stalled replication forks, DNA breaks and other chromosomal aberrations.

## Results

### PALB2-DBD is comprised of a low complexity amino acid sequence with predicted context dependent folding

PALB2-DBD sequence includes the N-terminal α-helix (aa 10–40) (46) which forms an antiparallel coiled-coil either with itself (referred to hereafter as PALB2-cc) to form a homodimer, or with the BRCA1 α-helix(47–49) (**Fig. 1A**). The remaining part is comprised of a low complexity sequence, typical for scaffold proteins, with several short low propensity α-helical regions (https://MobiDB.bio.unipd.it/Q86YC2 and https://alphafold.ebi.ac.uk/entry/Q86YC2) (50, 51). However, PALB2-DBD properties such as dimerization, localized DNA binding site, and complex strand exchange activity are not typically associated with disordered proteins. Moreover, the fragment is also relatively soluble under physiological conditions and is proteolysis-resistant during purification. This suggests that that oligomerization and/or DNA binding can stimulate the formation of additional α-helixes and folding. Charge-hydropathy plot (52, 53) generated using PONDR® server (www.pondr.com), places the sequence close to the folded/disordered boundary (**Fig. 1B**). The PALB2-DBD is a neutral peptide with a balanced content of hydrophobic, positively, and negatively charged amino acids distributed alone the entire sequence (**Fig. 1C, D**), as calculated using CIDER analysis (54), and characteristic for a context-dependent structural organization (54–58). Δ40-DBD sequence is placed at the boundary between “Janus sequences: Collapsed or expanded–context dependent” and “Weak polyampholytes & polyelectrolytes: Globule & tadpoles”. Analysis of this protein by the flDPnn platform (http://biomine.cs.vcu.edu/servers/flDPnn/), designed for the accurate intrinsic disorder prediction with putative propensities of disorder functions (59), further supports the excessive disorder-based interactivity of PALB2-DBD predicting multiple regions with a strong propensity for protein binding as well as DNA and RNA interactions.

PALB2-DBD sequence analysis using RIDAO server (https://ridao.app/) (60) which utilizes several predictors (mean disorder profile (MDP) error; PONDRs VLXT, VL3 and VSL2, IUPpred-Long, IUPred-Short and PONDR-FIT) suggest a mostly disordered structure (**Fig. 1E**), while multiple segments of the domain (aa 5-46, 49-58, 70-79, 86-100, 118-129, 135-143, 145-167, and 174-194) were identified by FuzDrop (https://fuzdrop.bio.unipd.it/predictor) (61, 62) as regions with context-dependent interactions (**Fig. 1F**). Furthermore, the PALB2-DBD is predicted to contain six disorder-based protein-protein interaction sites (aa 18-35, 74-86, 98-106, 113-118, 154-174, and 193-206) known as molecular recognition features (MoRFs), which are IDRs undergoing binding-induced folding at interaction with specific partners (63–68) (**Supplementary Figs. S1A, B**). Thus, the sequence analysis does not rule out potential context-dependent folding which may be required for novel strand exchange mechanism. Therefore, we used several experimental approaches to characterize the PALB2-DBD structure.

### PALB2-DBD structure is highly flexible lacking stable secondary structure elements beyond the N-terminal α-helix with and without DNA

We utilized circular dichroism (CD) spectroscopy to analyze the content of secondary structure elements (**Fig. 2**). The PALB2-DBD spectra are characterized by peaks corresponding to α-helical (223 nm) and disordered (204 nm) structures (**Fig. 2A**, solid line). The spectrum of the Δ40-DBD fragment lacking the N-terminal α-helix does not display a peak corresponding to α-helix (**Fig. 2B**). Therefore, CD spectroscopy did not detect secondary structure elements beyond the N-terminal coiled-coil region in PALB2-DBD. To investigate whether additional secondary structure elements can be stabilized by interactions with DNA, we measured CD spectra from PALB2-DBD and Δ40-DBD in the presence of excess ssDNA (dT_50_) (**Figs. 2A, B**, dashed lines) at buffer conditions compatible with DNA binding (**Supplementary Fig. S2**).

**Fig. 2.**
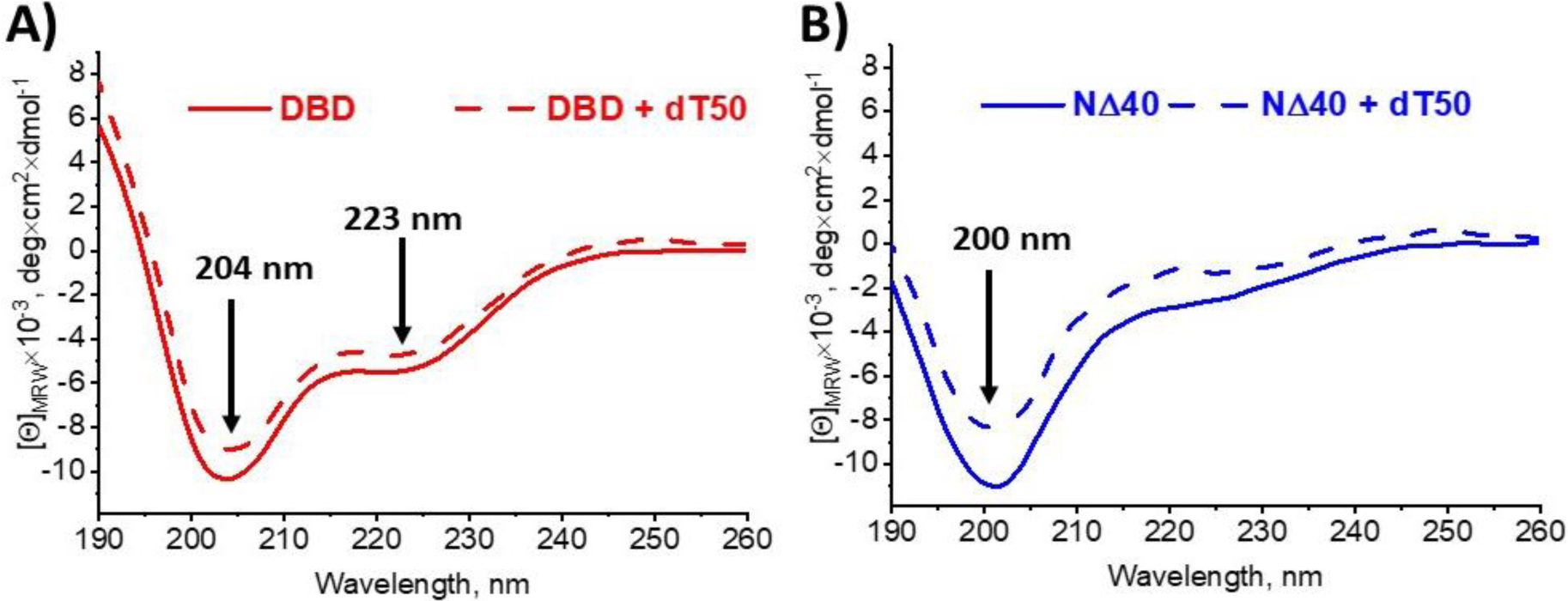
CD spectroscopy of PALB2-DBD. CD spectra of PALB2-DBD **(A)**, and of Δ40-DBD **(B)**. Solid lines correspond to the protein alone at 30 μM and dashed to the protein in the presence of 30 μM dT_50_. Spectrum were obtained in a buffer 10 mM NaPO_4_, 150 mM NaF, 2 mM CHAPS.

The results showed that neither PLAB2-DBD nor Δ40-DBD spectra were altered by addition of ssDNA. While this qualitative analysis does not rule out the presence of minor and/or transitional secondary structure elements nor the potential formation of additional α-helixes upon coiled-coil–mediated oligomerization of PALB2-DBD, it suggest that Δ40-DBD lacks stable secondary structure elements and folding even upon DNA binding, despite a localized major DNA-binding site typical for globular proteins.

As IDRs are characterized by high flexibility, we analyzed the local flexibility of different regions using continuous wave electron paramagnetic resonance (cwEPR) spectroscopy (**Fig. 3A-C**), which is particularly informative for structural characterization of IDPs and their interactions (69, 70). Four cysteines in human PALB2-DBD were alternatively substituted by alanines to create single cysteine variants at positions 11, 57, 77, and 162 (**Fig. 3D**). Lack of cysteines did not alter DNA binding properties of PALB2-DBD (**Supplementary Fig S2**). Two additional cysteines were introduced in the vicinity of the major DNA-binding site (S136C, S145C). The nitroxide-containing spin label 1-oxyl-2,2,5,5-tetramethyl-d3-pyrroline-3-methyl methane thiosulfonate (MTSSL) was covalently attached to the cysteine thiol group for EPR analysis. The EPR spectrum was measured for each mutant alone (**Fig. 3A**) and in the presence of ssDNA (**Fig. 3B, C**). Most spectra correspond to a highly flexible conformation of local peptide in the label vicinity. Values of the h(+1)/h(0) ratio correspond to those for other IDPs (**Fig. 3C**) (70), with the exception of position 11 at the beginning of the PALB2-cc where a lower value corresponds to a more rigid structure near the coiled-coil region. The overall tumbling rate of the complex (dimer molecular weight of 47 kDa) is too slow to affect the EPR spectrum. The antiparallel coiled-coil structure was previously detected for isolated α-helixes by NMR (46). To confirm that this interaction is preserved within the entire PALB2-DBD, we used DEER spectroscopy of protein labeled at position 11 (**Fig. 3E, F**). The measured distance of 47.3 Å is in good agreement with theoretical distribution calculated using NMR structure (**Fig. 3E**, insert).

**Fig. 3.**
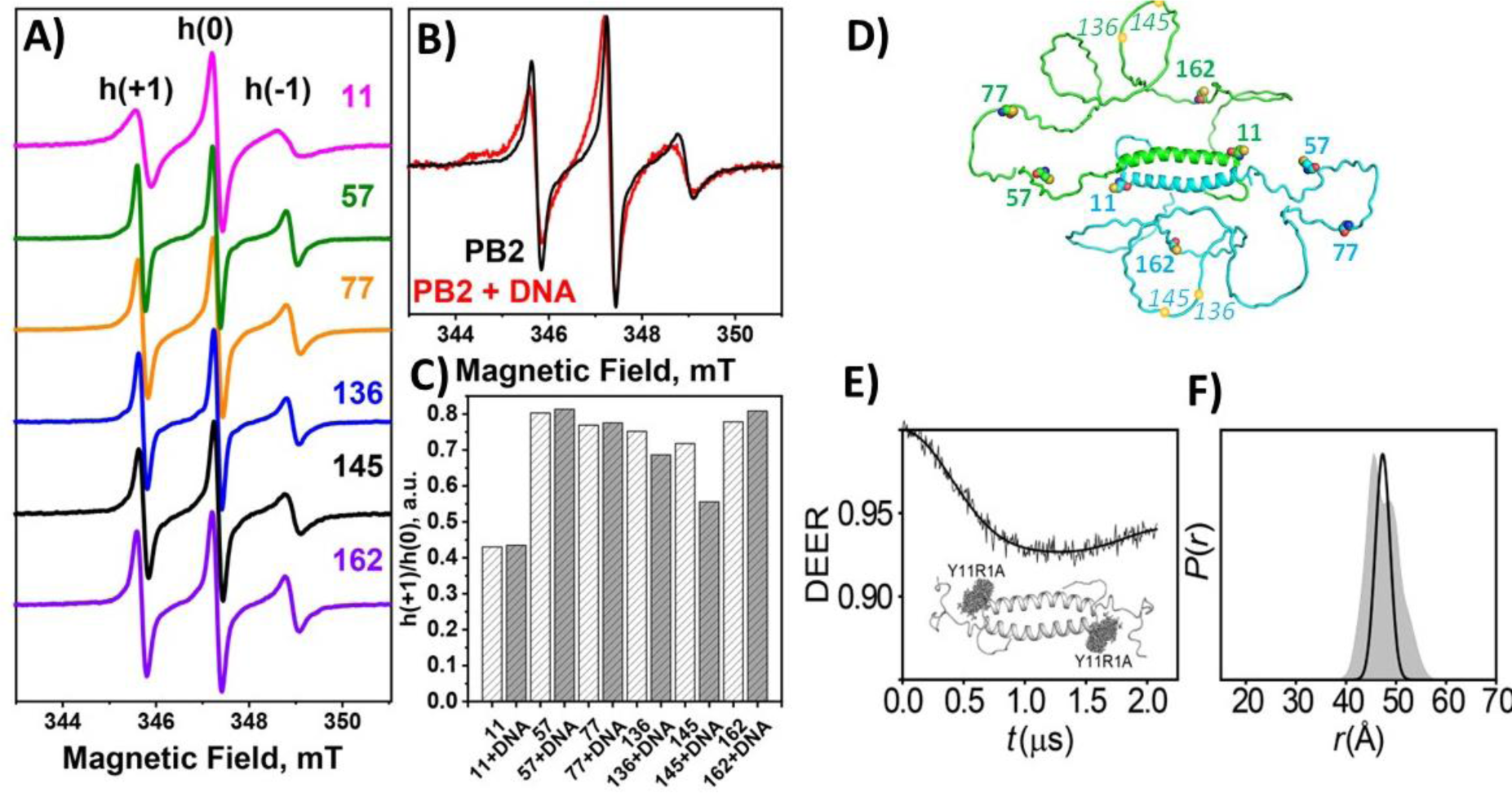
Continuous wave EPR and DEER spectroscopy of PALB2-DBD. **(A)** cwEPR spectra of PALB2-DBD labeled at different positions indicated by the number on the right. obtained for the **(B)** Overlay of PALB2-DBD spectra labeled at position 145 in free (black) and DNA-bound (red) states. **(C)** h(+1)/h(0) values for the labeled positions. Dark grey bars correspond to the signal in the presence of DNA. cwEPR data were obtained with 30 μM protein and 60 μM dT_40_ in buffer with 20 mM Tris-acetate pH 7, 100 mM NaCl, 10% DMSO, and 5% glycerol. **(D)** Hypothetical conformation of PALB2-DBD dimer generated by the AlphaFold program. Monomers are shown in cyan and green. Cysteine positions used for labeling are marked with spheres and aa numbers. **(E)** Raw DEER decay for the PALB2-DBD labeled at position 11 with fitting for the experimentally determined *P(r)*. **(F)** The corresponding *P*(*r)* distribution. The distribution predicted from the NMR structure (insert in (**E**)) is shaded gray.

The addition of dT_40_ causes a notable change in flexibility only at position 145 immediately next to the major DNA-binding site (146–149 aa) and a minor decrease at position 136 proximal to DNA binding site (**Fig. 3B, C**). Both regions are still more flexible even in the presence of DNA than at position 11. There was no change in local flexibility of the region at other positions (**Fig. 3C**). Thus, DNA binding does not affect structural flexibility beyond the immediate vicinity of the major DNA-binding site. High flexibility of the probed regions further confirms a lack of secondary structures formed inside the Δ40-DBD sequence, even within the oligomeric PALB2-DBD and in the DNA-bound state. The combined results indicate that PALB2-DBD has an intrinsically disordered and highly flexible structure beyond the N-terminal α-helix.

### A dimerization-dependent compaction of PALB2-DBD

IDPs have different functionally relevant structural organizations ranging from compact molten globule (MG) structures to extended random coil (RC) conformations, which can be distinguished by a hydrodynamic volume (*V_h_*) that is only 2–3 times larger than the volume of a folded globular structure of the same molecular weight for MG and up to 10 times larger for RC (71). Since the PALB2-DBD structural organization is further complicated by oligomerization (**Figs. 3E,F**, data below and reference (46)), we initially evaluate the compaction of monomeric Δ40-DBD using size exclusion chromatography (SEC) to compare the elution time of the protein in a DNA-binding buffer and in the presence of the strong denaturant guanidinium hydrochloride (GdmHCl), which forces RC formation (**Supplementary Fig. S3**). The protein elutes significantly earlier in 6 M GdmHCl than in non-denaturing DNA-binding buffer. The elution volume under nondenaturing conditions corresponds to a globular protein with a molecular weight only twice that of Δ40-DBD.

Next, we characterized structural organization of the PALB2-DBD dimer using small angle X-ray scattering (SAXS). SAXS can directly measure structural features of proteins in solution and is particularly valuable for IDPs that are not amenable to other structural techniques (72–74). SAXS measurements were conducted in-line with SEC and multiangle light scattering (MALS) to separate and identify different molecular species. We first analyzed PALB2-DBD in two different buffer conditions with 0.16 M or 0.5 M NaCl (**Supplementary Tables S1, S2 and Supplementary Figs S4, S5**), followed by studies on the L24A mutant and of the PALB2-DBD complex with dT_50_ (**Supplementary Tables S3, S4 and Supplementary Figs S6-8**). Two salt concentrations were initially used because PALB2-DBD is more soluble and less prone to aggregation in high salt buffer (0.5 M NaCl), whereas DNA binding is strongly inhibited by high salt concentrations (32). At 0.16 M NaCl PALB2-DBD retains DNA binding activity at submicromolar range (**Supplementary Fig. S2)**. MALS measurements unequivocally confirm the dimeric form of PALB2-DBD under both buffer conditions (**Fig. 4A, B**) and the monomeric form of L24A mutant at 0.16 M NaCl (**Fig. 4C**). Interestingly, previous measurements of the coiled-coil dimerization constant by NMR revealed *K_d_*=80 μM for the isolated N-terminal α-helical fragments (46). Retaining of the dimeric state by PALB2-DBD at lower concentrations even under nonequilibrium SEC conditions suggests that an interaction between disordered regions is synergistic with the coiled-coil interaction.

**Fig. 4.**
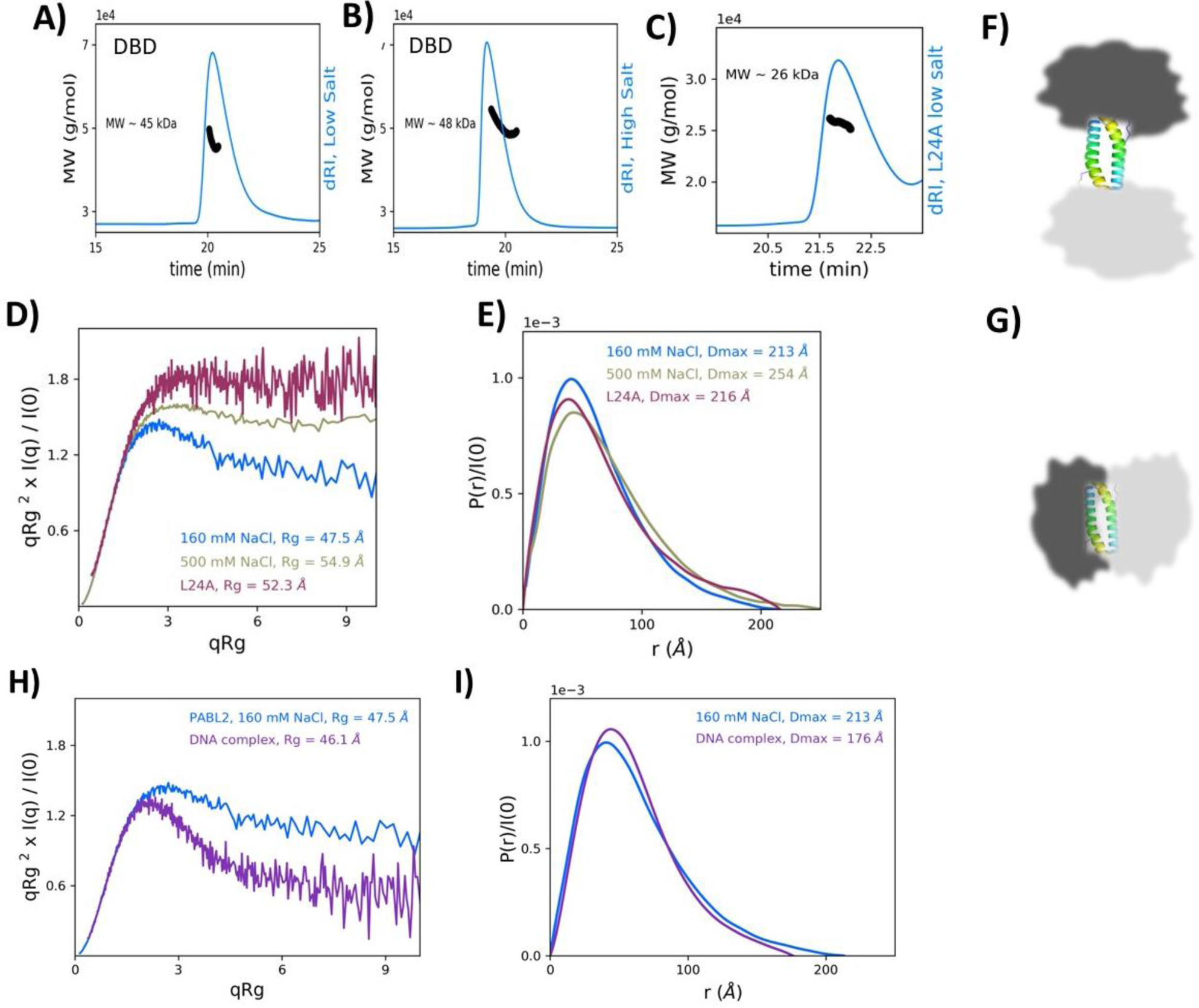
SAXS/MALS analysis of PALB2-DBD. **(A-C) SEC** elution profile (blue) measured by refractometer and MW estimation by MALS (black) of PALB2-DBD at 160 mM (**A**) and 500 mM (**B**) NaCl buffers and of L24A mutant (**C**) at 160 mM NaCl. **(D)** SAXS data for PALB2-DBD at high (green) and low (blue) NaCl concentration buffers and of L24A mutant at 160 mM NaCl (dark red), visualized as a dimensionless Kratky plot. Data were binned logarithmically by factor of 3 for visual clarity. **(E)** Pairwise distance distribution functions [*P*(*r*)] from SAXS measurements of same samples as in (D). **(F-G)** Two hypothetical models of IDRs attached to opposite ends of coiled-coil linker. A single domain length in *P*(*r*) distribution (E) suggests a compact dimer conformation (bottom). (**H)** SAXS data for PALB2-DBD (blue) and complex of PALB2-DBD with dT_50_ (magenta) visualized as a dimensionless Kratky plot. Data were binned logarithmically by factor of 3 for visual clarity. The SAXS profile of the DNA complex is significantly more compact, more closely resembling a scattering for a globular protein than for an IDP. **(I)** Pairwise distance distribution functions [*P*(*r*)] from SAXS measurements of same fragments as in (H).

SAXS analyses resulted in typical IDP scattering patterns, as illustrated by the slow decay of the tail when data are plotted as a dimensionless Kratky plot distribution (**Fig. 4D**). Distribution of PALB2-DBD at 500 mM NaCl appears slightly more compact than a typical disordered IDP, with the slight bell-shaped peak at ∼2.5 qR_g_ and flat tail (**Fig. 4D, teal**). These features are even more pronounced at low salt suggesting an additional compaction at low salt (**Fig. 4D, blue**). Likewise, the estimated values of *R_g_* are 54.9 Å and 47.5 Å for high and low salt conditions, correspondingly. Fitting the 0.5 M NaCl data to a molecular form factor (MFF) fitting protocol developed specifically for IDPs (75, 76) yields a Flory exponent ν=0.52, which describes a polymer dimension dependence on its length (*R_g_*∼N^ν^) (**Supplementary Data Table S2**). This value of ν is slightly lower than those for typical IDPs in good solvent (∼0.54–0.55) (75, 76), and suggests a relatively compact overall conformation, assuming that this system behaves similarly to a single-chain IDP with a total length of 400 aa due to the almost end-to-end dimerization. In fact, these compaction parameters at 0.5 M NaCl are close to the corresponding values reported for an unstructured PNt domain with smaller single chain size (335 aa) (75). The *R_g_* (47.5 Å) and ν (0.47) values are notably lower in the low salt condition than in the high salt condition, suggesting a role for polar interactions in structure compaction. Interestingly, the distribution obtained for L24A monomer suggests a more disordered state (**Fig. 4D, dark red**) than the dimer under either salt condition. In fact, an estimated *R_g_* value of 52.3 Å (ν=0.54) of the monomeric L24A mutant even larger than the *R_g_* of the dimer under same conditions (**Figs. 4D, E**), suggesting that the interaction between IDRs subunits leads to compaction of each monomer and of the entire dimer.

Two IDRs in the PALB2-DBD dimer are located at opposite ends of 40 Å-long antiparallel coiled-coil structure and can form two independent or spatially distinct MGs. However, only one dominant length scale is observed in the pairwise distance distribution [*P*(*r*)] measured in SAXS (**Fig. 4E**) supporting a compact structure (**Fig. 4G**), ruling out the dumbbell model with two distinct MGs (**Fig. 4F**). While the distinction between two models may be less pronounced in case of the disordered structure, the relatively small *R_g_* values from SAXS experiments with the dimer also indicates a compact overall organization. These combined data suggest that the monomer adopts a typical MG-like IDP structure stabilized by intramolecular interactions and similar interactions between two IDRs contribute to synergistic dimerization with a coiled-coil interaction, thereby leading to an additional structural compaction of the entire dimer.

### Compaction of PALB2-DBD upon DNA binding

Following studies on the protein alone, we conducted SEC-MALS-SAXS data on PALB2-DBD with dT_50_ DNA in a low salt buffer with 160 mM NaCl. As this experiment was performed at a different facility (Cornell High Energy Synchrotron Source, CHESS), we also collected control data for PALB2-DBD alone in 160 mM NaCl buffer. The scattering profile for PALB2-DBD alone is in excellent agreement with the previous APS experiment (**Supplementary Fig. S9**), albeit with slightly larger estimated *R_g_* and *P(r) D_max_* values, likely as a result of increased propensity for sample radiation damage at CHESS due to differences in beamline infrastructure (77). For data collected at CHESS, deconvolution by EFA (78) and/or REGALS (79) was necessary to separate out the component of interest from other species and accumulated radiation damage (**Supplementary Figs S10-S12**). Addition of DNA to protein resulted in a significantly more compact structure of the dimer, with a Kratky plot shape more characteristic of a globular protein than an IDP (**Fig. 4H**). Correspondingly, the *R_g_* of the complex with DNA is 46.1 Å, smaller than PALB2-DBD alone under identical experimental conditions (47.5 Å) despite being in complex with DNA. SEC elution time supports the dimeric form of the complex with dT_50_; however, MALS data were highly variable and inconclusive.

### SAXS-directed modeling of structural ensembles

To obtain further insight into PALB2-DBD structural organization using experimental SAXS results, we performed molecular structure modeling of the protein dimer and of L24A mutant monomer using the program SASSIE (80) (**Supplementary Table S5**). An initial pool of structures were generated using the Complex Monte-Carlo algorithm in SASSIE with fixed N-terminal α-helixes and a dimeric coiled-coil interface (aa 11–40) as determined by NMR (46). The starting model of a dimer was generated by the AlphaFold program using the ColabFold Google server (81) (**Fig. 3D**). The resulting trajectory yielded 34,919 structures, from which a subensemble of structures (selected using GAJOE (73)) provided a good fit (χ^2^=1.01) to the experimental data of the system in 0.5 M NaCl buffer (**Fig. 5A**). The poll of the best-fit ensemble is bimodal with the majority with lower *R_g_* ∼45 Å and a second population with *R_g_* ∼62 Å (**Fig. 5B**). Although the experimental Guinier *R_g_*=54.9 Å is significantly lower than the average *R_g_* ∼71 Å of the entire pool of all randomly generated structures, enough compact structures were sampled that the average *R_g_* of the selected ensemble (56.7 Å) is similar to the experimental value.

**Fig. 5.**
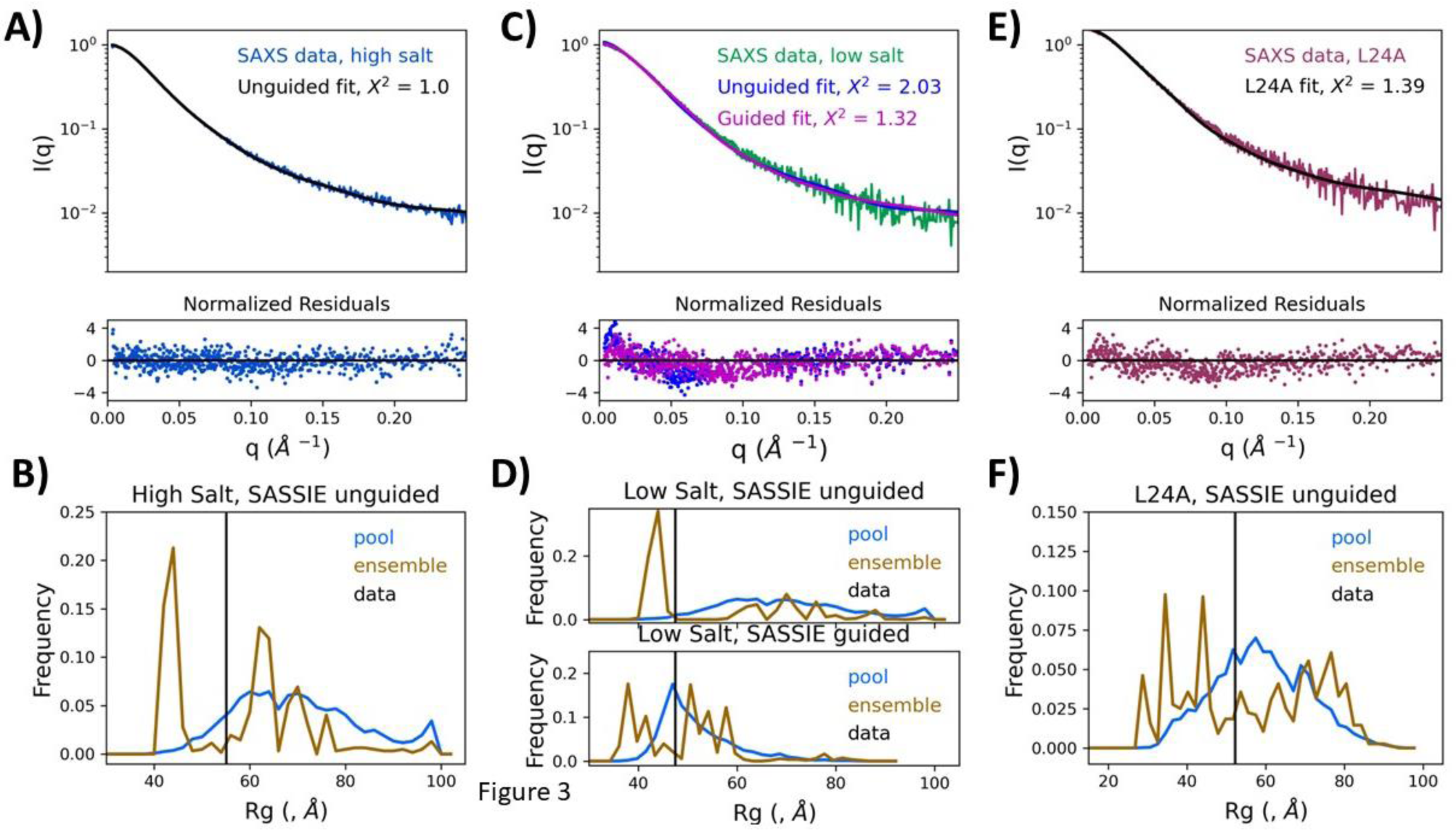
SASSIE fits to experimental SAXS data. **(A)** Fit of SAXS-derived experimental distance distribution of PALB2-DBD in 500 mM NaCl with theoretical distribution of best fit models calculated from unguided SASSIE models using GAJOE; **(B)** *R_g_* distribution of the entire pool of 34,919 structures (blue), experimental *R_g_* (black bar), and distribution of *R_g_* of the best-fit pool structures (brown); **(C-D)** similar plots to that of in (A,B) with both unguided (blue) and guided (red) fits shown in (C) with corresponding *R_g_* distributions shown in top and bottom panels in (D) for unguided and guided fits, correspondingly; **(E-F)** similar fitting and *R_g_* distributions as in (A,B) for data from L24A monomeric mutant in 160 mM NaCl buffer.

Fitting data obtained at the low salt concentration (0.16 M NaCl) was more challenging, likely due to the low population of compact structures generated by the unguided SASSIE Monte Carlo simulation. Fitting the high salt condition data using the same structural pool generated for fitting the low salt condition resulted a poorer fit (χ^2^=2.03, **Fig. 5C)**. Guiding the simulation trajectory to an average *R_g_* (the experimental *R_g_* of 47.5 Å) during modeling resulted in an improved fit (χ^2^=1.32), as simulation was successfully able to generate a structural pool with a much smaller average *R_g_* (visualized in the blue distributions in Fig. **5D**). A similar bimodal distribution of theoretical *R_g_* for the selected ensemble was obtained even with guided modeling (**Fig. 5D**).

We also performed similar molecular modeling to the L24A SAXS data. An initial ensemble of 38,042 structures was generated using the Monomer Monte Carlo simulation implemented in SASSIE. The starting model used was chain A from the AlphaFold-generated dimer, as described above, with the L24A mutation performed using the simple mutagenesis tool in Pymol. The initial pool of generated structures was relatively well distributed around the experimental *R_g_* (**Fig. 5F, blue**), and a final sub-ensemble (selected using GAJOE (73)) fit the experimental data well (χ^2^=1.39). The distribution of the final ensemble was again somewhat bimodal (**Fig. 5F, red**), though less pronounced than for the dimer.

### Single-molecule FRET analysis reveals that PALB2-DBD dynamically condenses DNA

Two studied strand exchange protein families, RecA and Rad52, are characterized by significantly different DNA biding mechanisms. RecA-like recombinases form nucleoprotein filaments stretching out ssDNA (82–84). Rad52 wraps ssDNA around a toroidal oligomeric structure (24, 25, 85). Our previous intensity-based Förster resonance energy transfer (FRET) experiment suggested a Rad52-like mechanism where protein interaction led to increased FRET between Cy3 and Cy5 dyes located at the ends of doubly labeled dT_40_ or dT_70_ oligonucleotides (32). Titration by PALB2-DBD led to higher FRET efficiency, thus resembling the results of a similar experiment with Rad52 (85). However, these measurements were performed on an ensemble averaged in a fluorimeter, so the molecular species and sequence of events that generated the result could not be precisely identified, particularly considering the disordered nature of PALB2-DBD and oligomerization.

To address the limitation of our previous studies, we performed comparable experiments utilizing single-molecule FRET (smFRET). We used a confocal setup to determine the FRET efficiency of freely diffusing single DNA molecules labeled with the FRET pair Cy3/Cy5 in the absence and presence of PALB2-DBD. We used Pulsed interleaved excitation (PIE) to calculate the stoichiometry for each molecule, as the PIE value enables the identification of molecules containing the correct 1:1 ratio of donor and acceptor (**Fig. 6A**) and the detection of other photophysical phenomena such as bleaching, blinking, and protein-induced fluorescent enhancement (PIFE) (86–89). Measurements were performed with several ssDNA substrates with fluorescent dyes placed at different distances: dT_50_ with Cy3 at the 5′ end and Cy5 at positions 25 or 50, and dT_70_ labeled at the 5′ and 3′ ends. Cy3-dT_70_-Cy5 and Cy3-dT_50_-Cy5 substrates in solution show a single, symmetric distribution with mean FRET of 0.11 (**Fig. 6A**) and 0.18 (**Fig. 6B**), respectively, in agreement with published results (90). The addition of PALB2-DBD significantly increased mean FRET, thereby confirming observations from our previous studies(32). Two distinct FRET populations were observed in the presence of PALB2, corresponding to two different states of DNA bound to the protein, one significantly more condensed than the other with mean FRET of 0.34 and 0.71 in case of Cy3-dT_70_-Cy5 (**Fig. 6A**, the grey panel above each 2D plot). The mean FRET value for each peak is invariant for a given DNA fragment, whereas the population of each state changes upon titration by PALB2-DBD. A medium FRET state of Cy3-dT_70_-Cy5 is a predominant species at 1 μM PALB2-DBD, whereas high FRET is a major peak at 3 μM PALB2-DBD (**Fig. 6A**). Titration of dT_50_ by PALB2-DBD (**Fig. 5B**) revealed similar distributions of two populations of invariant mean FRET peaks of 0.45 and 0.79, with a high FRET peak becoming more predominant at high protein concentration. These results suggest that PALB2-DBD binds to multiple sites within the same DNA strand, thereby dynamically controlling its conformation. They also suggest that the PALB2-DBD concentration is an important factor in controlling DNA conformation.

**Fig. 6.**
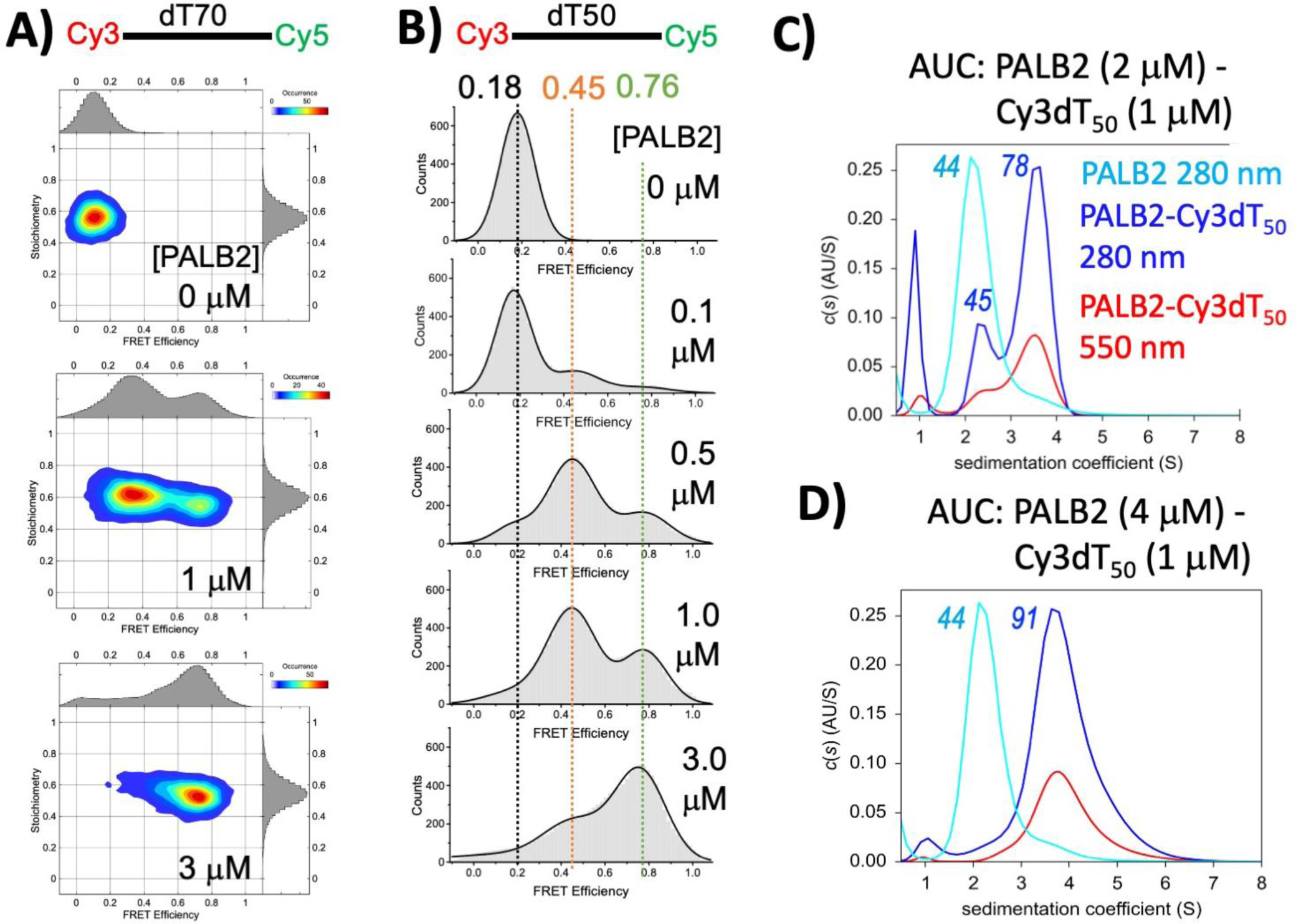
Bimodal DNA compaction revealed by confocal smFRET and two oligomeric states revealed by AUC. **(A)** 2D plots of FRET efficiency versus stoichiometry for 100 pM Cy3-dT_70_-Cy5 alone (top panel) and in the presence of 1.0 μM (middle panel) and 3.0 μM PALB2-DBD (bottom panel) in a buffer with 160 mM NaCl, 20 mM HEPES pH 7.5, 1 mM TCEP. FRET efficiency histograms are shown in grey above each plot and FRET pair stoichiometry in grey on the right side of each plot. **(B)** FRET histograms (grey) of 100 pM Cy3-dT_50_-Cy5 titrated by PALB2-DBD. Black lines show fitting of data using three species as Gaussian distributions. Protein concentrations are shown on each graph. Mean FRET values of major peaks are shown with color corresponding to each peak. **(C)** Sedimentation coefficient distribution of PALB2-DBD at 6 μM (cyan), of PALB2-DBD at 2 μM in the presence of 1 μM Cy3-dT_50_ (blue, red) measured at absorption wavelengths of 280 nm (blue) and at 550 nm (red) in identical buffer to that of in (A,B). **(D)** Similar distributions as in (C) but at 4 μM PALB2-DBD mixed with 1 μM Cy3-dT_50_.

A similar trend was observed for a dT_50_ substrate with Cy3 and Cy5 labels placed 25 nucleotides apart (**Supplementary Fig. S13A**). Mean FRET for free DNA is 0.45, the major peak at 1 μM PALB2-DBD is at FRET of 0.71, and the peak for 3 μM PALB2-DBD is at FRET of 0.89. DNA compaction is significant for this substrate as well, although it is challenging to deconvolute two separate peaks at each protein concentration due to the initially high FRET for free DNA. A similar trend of DNA compaction proportional to the number of nucleotides between labels is observed for all three cases. For example, changes in FRET between the peak of free DNA and the first peak of DNA in complex with PALB2-DBD for distances of 70, 50, and 25 nucleotides were from 0.11 to 0.34, from 0.18 to 0.45, and from 0.5 to 0.7, respectively. These results ruled out a Rad52-like wrapping model of DNA binding, in which the FRET change between remote ends is greater than that between labels situated closer in the nucleotide sequence.

The broad distributions of each peak suggest a random coil model of ssDNA conformation, in which the average distance between ends is restricted due to interactions with two major DNA-binding sites in a compact PALB2-DBD dimer. In this model, two random parts of ssDNA should interact with two DNA-binding sites situated within an average *R_g_* distance from each other **(Fig. 7**). This model explains an invariant position of the peak with an intermediate mean FRET. The mechanism of an additional DNA compaction corresponding to a high FRET peak remains to be further investigated. One hypothesis is that an additional compaction is caused by a higher oligomeric state of PALB2-DBD. To evaluate the role of PALB2-DBD oligomerization in DNA compaction, we tested a Δ40-DBD variant lacking the coiled-coil dimerization interface using Cy3-dT_50_-Cy5 (**Supplementary Fig. S13B**). DNA compaction also was observed in this case, albeit at higher protein concentrations, reflecting a significantly reduced DNA binding affinity (32). There was a fraction of free DNA in solution even at 10 μM protein. FRET distributions were significantly broader and more heterogeneous corresponding to multiple species of DNA conformations. The lack of two distinct peaks in the case of Δ40-DBD indirectly supports oligomerization-dependent DNA compaction. In this case, the Δ40-DBD will form oligomers by addition of a single monomer (dimers, trimers, tetramers) resulting in a broad distribution of DNA conformations, while PALB2-DBD forms dimers and, potentially, tetramers resulting in a more distinct FRET values.

**Fig. 7.**
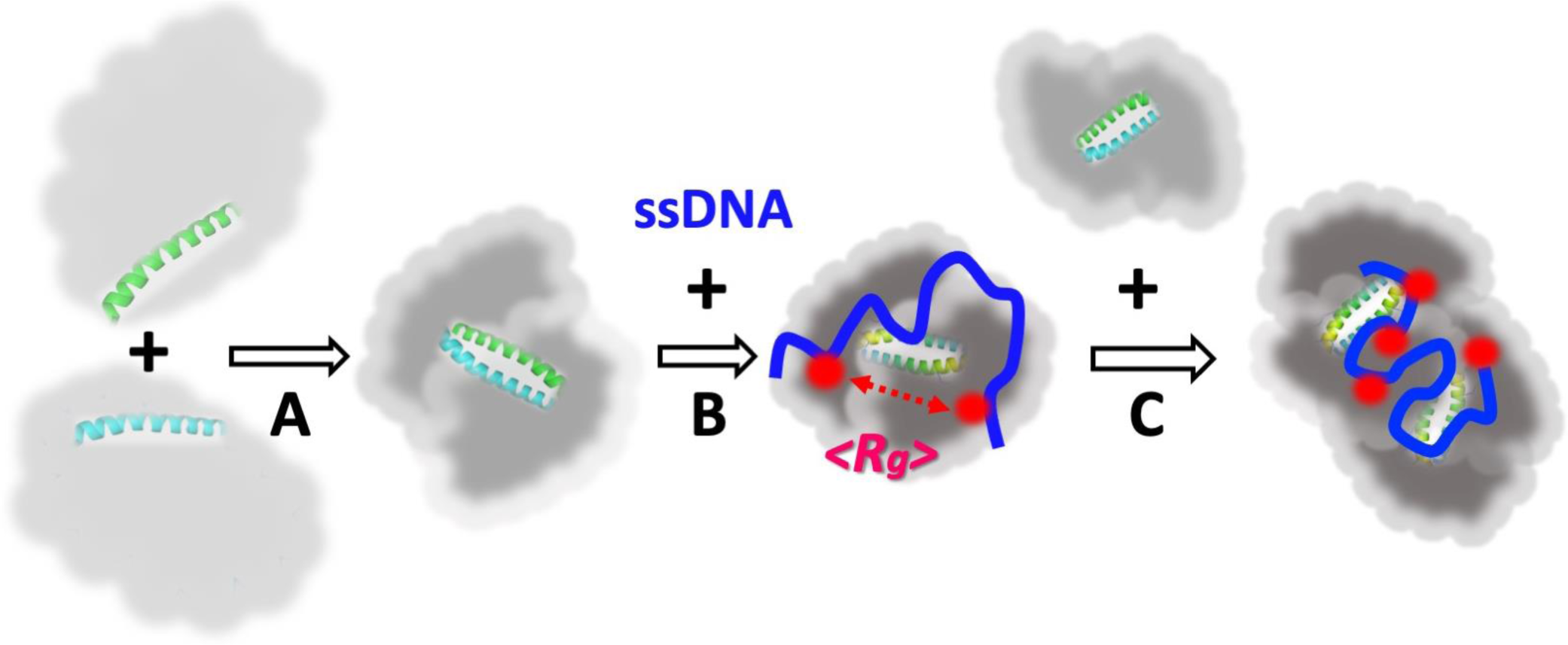
Model of oligomerization-dependent compaction of PALB2-DBD and of ssDNA compaction by PALB2-DBD dimer and tetramer. **(A)** PALB2-DBD dimerization is mediated by coiled-coil and IDR-IDR interactions leading to compaction of the structure. **B)** ssDNA (blue) binds to PALB2-DBD dimer in a random but a significantly more compact conformation, comparatively to its free state in solution, due to interaction with two major DNA-binding sites (red) situated at an average distance of *R_g_* from each other. Interaction with ssDNA leads to an additional compaction of PALB2-DBD dimer. **(C)** Higher concentration of PALB2-DBD leads to tetramerization and additional compaction of ssDNA bound to tetramers.

The tetramerization hypothesis is supported by a previously observed 1:4 stoichiometry of DNA binding in fluorescence polarization and ensemble FRET assays (32), but is contradicted by a dimeric state of PALB2-DBD measured in SEC/MALS/SAXS experiments. We conducted sedimentation velocity experiments with PALB2-DBD under identical conditions in the presence of Cy3-dT_50_ (**Fig. 6C, D**). DNA was labeled with Cy3 to distinguish its position from that of protein-only. Measurements were repeated at two different protein concentrations corresponding to DNA:protein molar ratios of 1:2 and 1:4 with DNA at 1 μM concentration in both experiments. PALB2-DBD at 6 μM sediments at molecular weight of ∼45 kDa corresponding to a dimer (cyan) at concentration of. Three peaks were observed upon addition of ssDNA at 1:2 DNA:protein ratio, one at low molecular weight position, corresponding to free DNA, the second at position of PALB2-DBD dimer (an apparent MW=45.2 kDa), and the third at position with molecular weight of ∼78 kDa, corresponding to the tetramer. A low apparent molecular weight (theoretical value is 92 kDa) of this peak is likely due to the dynamic exchange between different oligomeric states. All three peaks are present in absorption profile at 550 nm, corresponding to maximum absorption of Cy3, confirming that Cy3-dT_50_ is bound to dimers and tetramers. Only one, albeit wider peak corresponding to the tetramer is present at 1:4 protein ratio with a calculated molecular weight of 91 kDa. Note, that AUC experiment is conducted under equilibrium conditions while SEC is a non-equilibrium experiment where unstable oligomeric complexes dissociate during migration through the column. This can explain the lack of tetrameric species observed in SEC/MALS/SAXS experiment. Assuming that high FRET value peak corresponds to tetramers observed in AUC both confirm the presence of tetramers in the presence of ssDNA. Since both dimers and tetramers are formed at conditions with large stoichiometric excess of protein versus ssDNA, we believe that tetramerization depends on total protein concentration.

## Discussion

Our data indicate that PALB2-DBD is an intrinsically disordered domain that forms a dimer synergistically stabilized by a coiled-coil interface between N-terminal α-helices and the interactions between IDRs. Although PALB2-DBD is comprised of a low complexity sequence, it was unexpected that an unstructured protein can support an elaborate multisubstrate and multistep strand exchange. Results render novel structural and DNA interaction properties of PALB2-DBD which have not been described for any other characterized DNA binding domains, to our knowledge. While the mechanism of strand exchange remains to be investigated, results clearly suggest a completely different mechanism from that of RecA-like recombinases and Rad52.

SEC and SAXS results revealed that the PALB2-DBD dimer has a compact structure not typical for scaffold structures interacting with multiple proteins. This compaction is achieved through favored intermolecular and intramolecular interactions between two IDRs as the monomeric mutant is significantly less compact than the dimer. The Flory scaling exponent describing a polymer dimension dependence on its length (ν) is around 0.5, which is expected for a non-self-avoiding random coil and corresponds to a Θ solvent where intermolecular and solvent interactions counterbalance each other (76). A noticeable reduction of *R_g_* and ν in a low salt buffer suggests contribution of polar intermolecular interactions to the compaction. The interaction between two IDRs is weak, since Δ40-DBD and L24A are monomeric in solution. However, they are synergistic with the coiled-coil interactions. A single dominant length scale observed in the pairwise distance distribution [*P*(*r*)] measured in SAXS (**Fig. 4E, I**) further supports interactions between entire domains and not only coiled-coil helixes. It is tempting to hypothesize that similar interaction between IDRs of other proteins can contribute to heterooligomerization within DNA repair complexes. For example, PALB2 binds BRCA1 through the same N-terminal α-helix that forms a coiled-coil interface with the BRCA1 α-helix (49, 91) surrounded IDRs (92–94). Thus, these interactions can be strengthened by attractions between IDRs of two proteins. Importantly, SAXS data demonstrated that ssDNA binding leads to additional compaction of the dimer. The mechanism of such compaction has to be further investigated through computational modeling and mutagenesis studies.

We observed significant compaction of ssDNA on PALB2-DBD binding, ruling out a RecA-like mechanism, since RecA-like recombinases stretch ssDNA (95). In case of Rad52, ssDNA wraps around the toroidal oligomer backbone, bringing distant ends of DNA in close proximity (85, 96). However, the wrapping model does not fit behavior ssDNA bound to PALB2 since changes of FRET efficiency are proportional to the number of nucleotides between labels, while in a wrapping model change between remote ends are greater than between more close positions (e.g. full circle versus half circle wrapping). Thus, the mechanism of PALB2-DBD interaction with ssDNA differs from those of RecA and Rad52.

Wide FRET distributions of PALB2-DBD–bound ssDNA and the invariant positions of observed peaks suggest random coil folding with an imposed distance restriction. The interaction of ssDNA at two random positions with two DNA-binding sites of the dimer (and potentially with minor DNA-binding sites) situated within an average *R_g_* distance from each other should lead to the observed compaction. The results indicate approximately two-fold DNA compaction, with a medium FRET value of Cy3-dT_50_-Cy5 bound to PALB2-DBD equal to that of unbound DNA with 25 nucleotides between Cy3 and Cy5. Importantly, the compaction of DNA directly correlates with the compaction of the dimer upon DNA binding.

The mechanism behind the formation of high FRET peaks for each substrate is less clear. We hypothesize that this conformation can be formed by interaction with tetrameric PALB2-DBD, as previously suggested and observed in AUC experiment. Similar attraction forces between IDRs, which stabilize the dimer, can synergistically stimulate interactions between two dimers along with DNA binding. Unlike folded protein with a defined protein-interaction interface supporting stable interaction, IDRs form a dynamic ensemble of multiple conformations where a specific sequence region can be involved in multiple dynamic interactions. Additional compaction of DNA bound to a larger complex can be achieved by condensation of DNA-binding sites around a ssDNA molecule. The physiological role of PALB2-DBD tetramerization remains unclear considering the required stoichiometric excess or relatively high concentration of protein, which may not be achieved for a full-length protein and/or its functional complexes with even larger BRCA2 and BRCA1. However, attractions between IDRs can stimulate formation of heterooligomeric complexes with disordered parts of BRCA1/BRCA2 and other protein partners or the formation of DNA-repair condensates through liquid-liquid phase separation (LLPS). The probability of forming a droplet state through LLPS (p_LLPS_) for PALB2-DBD and PALB2 is very high, as calculated using FuzDrop analysis (0.9622 and 0.9412, respectively, **Supplementary Fig. S14**). Since these values significantly exceed the 0.6 threshold, both PALB2-DBD and PALB2 are considered as potential droplet-drivers, which can spontaneously undergo liquid-liquid phase separation (61). Therefore, the observed limited oligomerization of PALB2-DBD and multivalent interactions (coiled-coil, IDR-IDR, protein-DNA) may reflect its ability to stimulate formation of biocondensates under certain conditions, which remained to be identified.

Previously reported strand annealing and strand exchange properties of PALB2-DBD combined with a dynamic nature of its structure and DNA interaction suggest a potential chaperone-like mechanism where transient iterative interactions can promote strand separation and reannealing. Chaperon activities are critical for RNA metabolism due to the complexity, diversity, and the dynamic nature of RNA structures and their complexes. Various mechanisms are supported by a wide spectrum of proteins ranging from ATP-dependent helicases to small IDPs (97, 98). *Escherichia coli* StpA protein promotes RNA strand annealing and strand displacement (99), similar to those reported for PALB2-DBD (32). SptA is a small globular protein that forms dimers required for these activities. Other examples include the structured globular FANCA protein of the Fanconi anemia (FA) complex (28) and limited reports of D-loop formation by FET proteins critical for RNA metabolism, specifically pro-oncoprotein TLS/FUS and the human splicing factor PSF/hPOMp100 (29–31). The latter proteins include structured domains (RNA recognition motif RRM and zinc finger), sequences of low complexity, and prion-like RGG domains implicated in higher-order self-assemblies. There are examples of disordered RNA chaperones composed of positively charged peptides that promote the formation of compact nucleic acid conformations by acting as polypeptide counterions to overcome repulsive electrostatic interactions of RNA or DNA chains (43–45). The mechanism of DNA compaction by PALB2 differs from such model as PALB2-DBD has overall neutral charge and significant hydrophobicity. Chaperone activities are not implicated in DNA metabolism due to significantly more restricted conformational space and size of chromosomal fragments. However, multiple DNA metabolism processes (e.g., repair of replication forks collided with transcription complexes and replication fork reversal) require the formation of transient multichain DNA and RNA-DNA structures and other higher-order nucleic acid interactions (100–102) that may benefit from strand exchange activity of PALB2-DBD or of similar unstructured domains of DNA-repair proteins. The fact that PALB2 resides at the promoter regions of highly transcribed genes under nondamaging conditions (103) further supports this functional hypothesis. Moreover, DNA-binding proteins are enriched in disordered domains/regions, have higher disordered content in eukaryotes (104), and many IDRs coincide with DNA-binding domains (92). Such regions may simply increase the chromatin affinity by transient DNA binding without forming a strong roadblock on DNA, except in cases of structure-specific interactions like nucleolin-G–quadruplex recognition (105). It is tempting to suggest that DNA-binding IDRs may have evolved to perform more complex reactions, such as a described strand exchange or DNA chaperon activities, using an evolutionarily inexpensive functional unit that facilitates formation and resolution of transient multichain DNA/RNA intermediates during chromatin replication and repair. Although this proposed reaction is more complex than others reported for IDRs, it depends on simple structural requirements and similar IDRs can be incorporated into larger proteins (e.g., DNA-binding regions in scaffold BRCA1 (92–94) and BRCA2 (106, 107)). Analysis of DNA-binding domain of BRCA1 show rather impressive overall similarity to the similar profiles generated for PALB2-DBD (**Supplementary Fig S15**) providing support to the validity of this hypothesis. These disordered domains can function synergistically with RAD51 (32, 40, 108), or they can independently perform a required reaction without forming stable presynaptic RAD51 filaments. Furthermore, such IDRs often have high propensity for LLPS formation, suggesting that they can stimulate formation of DNA repair condensates and function within such condensates. For example, sequence analysis of BRCA1 DBD region in vicinity of PALB2-binding site revealed a high probability of droplet formation by this region (**Supplementary Fig S15E**). This is a very compelling possibility, as the LLPS-driven formation of various membrane-less organelles (MLOs) and biomolecular condensates is considered now as one of the crucial organizing principles of the intracellular space responsible for regulation and control of numerous cellular processes (109). Among these MLOs related to the subject of this study are the DNA repair foci, formation of which represents a part of the DNA damage response (110–114); Promyelocytic leukemia Nuclear Bodies (PML NBs) that are physically associated with chromatin (115) and related to the chromatin function; liquid condensates on DNA/RNA matrices containing related proteins and G-quadruplexes (G4s) and responsible for reparation, transcription, genome integrity maintenance, and chromatin remodeling (116); nuclear condensates containing polycomb group (PcG) proteins that contribute to the reshaping of chromatin architecture under the physiological and pathological conditions (117); and intrinsic chromatin condensates related to various chromatin-centric processes, such as transcription, loop extrusion, and remodeling (118) to name a few. Therefore, it is likely that the LLPS-driven formation of biomolecular condensates containing this protein and nucleic acids can play a role in complex functionality of PALB2-DBD and similar regions of other DNA-binding IDRs,

## Materials and Methods

### Cloning and purification

PALB2 N-terminal fragments (1–195 I195W) was cloned into a pSMT-MBP plasmid containing an N-terminal 6×His-SUMO and a C-terminal MBP tag using Gibson assembly. The pSMT-MBP plasmid was created from a pET28b^+^–based pSMT3 plasmid (gift of Dr. R. A. Kovall, University of Cincinnati) by insertion of a TEV cleavage site and MBP at the ORF C-terminus. Mutations were introduced using the Stratagene QuikChange protocol. All constructs were fully sequenced and verified. Constructs were transformed into BL21* cells for expression. Cell cultures were grown in TB to OD_600_=1.6, and protein expression was induced by 0.2 mM IPTG overnight at 16°C. Cell suspensions were centrifuged at 4,000 rpm for 15 min. Cell pellets were resuspended in lysis buffer (25 mM HEPES pH 8.0, 1 M NaCl, 10% glycerol, 2 mM CHAPS, 1 mM TCEP, 0.3% Brij35, 2 mM EDTA, 1 mM PMSF), frozen in liquid nitrogen, and stored at −80°C. Frozen cell suspension was thawed and lysed with lysozyme at 0.25 mg/ml for 20 min at room temperature, followed by three rounds of sonication (50% output and 50% pulsar setting for 4 min). Cell debris was removed by centrifugation at 30,600 *g* for 45 min. Supernatant was loaded onto an amylose resin column (New England Biolabs) equilibrated with binding buffer (25 mM HEPES pH 8.0, 1 M NaCl, 10% glycerol, 2 mM CHAPS, 1 mM TCEP). Resin was washed with binding buffer containing 0.05% NP40 and 1 mM EDTA, followed by a second wash with low-salt buffer (50 mM NaCl, 25 mM HEPES pH 8.0, 1 mM TCEP, 2 mM CHAPS, 0.2% NP40, 1 mM EDTA). Protein was eluted in a binding buffer containing 20 mM maltose. SUMO and MBP tags were cleaved using Ulp1 and TEV proteases, respectively. Protein was diluted five-fold with binding buffer without NaCl to bring the NaCl concentration to 200 mM, loaded onto a Hi-Trap heparin affinity column (2×5 ml, GE Life Sciences), and eluted with a salt gradient (200–800 mM NaCl). Protein eluted from the column in the ∼600 mM NaCl fraction. Gel filtration was performed using a Superdex-200 10/300GL column (GE Life Sciences).

All DNA substrates were synthesized by Integrated DNA Technilogies (IDTDNA) Inc.

### Circular dichroism (CD)

A Zeba Spin desalting column (Thermo Scientific) was used to exchange protein buffer to buffer containing 10 mM NaPO_4_, 150 mM NaF, 2 mM CHAPS. Samples were centrifuged at 16,000 *g* to remove aggregates, and the final concentration was verified. Measurements were performed using a Jasco J-715. Four samples were used for CD measurements: (1) 30 µM PALB2-DBD, (2) 30 µM Δ40-DBD, (3) 30 µM PALB2-DBD with 30 µM dT_50_, and (4) 30 µM Δ40-DBD with 30 µM dT_50_. Each spectrum is an average of 10 scans.

### Electron paramagnetic resonance (EPR) analysis

Monocysteine PALB2 mutant proteins were labelled with MTSSL [S-(1-oxyl-2,2,5,5-tetramethyl-2,5-dihydro-1H-pyrrol-3-yl)methyl methanesulfonothioate] in a sample free of reducing agents. Unbound label was separated on a Superdex-200 10/300 gel filtration column. Spectra were measured using 30 µM protein in 20 mM Tris-acetate pH 7.0, 100 mM NaCl, 10% DMSO, and 5% glycerol with or without 60 µM dT_40_. Continuous wave EPR spectroscopy (cwEPR) was performed on an EMX spectrometer operating at X-band frequency (9.77 GHz) at room temperature. A dielectric resonator (ER4123D) and 25 µl borosilicate glass capillaries were used. The center field was set to 3474 G and the sweep width was 150 G. The modulation amplitude, microwave power, and resolution were set to 1.6 G, 2 mW, and 1125 points, respectively.

Distance measurements are obtained using a Bruker 580 pulsed EPR spectrometer operating at Q-band frequencies (34 GHz) using DEER with a standard four-pulse protocol as in previous publications(119–121). Glycerol is added to samples as a cryoprotectant, and DEER experiments are performed at 83° K. Primary DEER decay was analyzed using a home-written software operating in the Matlab (MathWorks) environment as previously described (122). Comparison of the experimental distance distribution with the NMR structure (46) using a rotamer library approach was facilitated by the MMM 2018.2 software package (123). Rotamer library calculation was conducted at 298 K.

### SEC-MALS-SAXS analysis

SAXS was performed at BioCAT (beamline 18ID at the Advanced Photon Source, Chicago) with in-line size exclusion chromatography (SEC) to separate sample from aggregates and other contaminants, thereby ensuring optimal sample quality. Multiangle light scattering (MALS), dynamic light scattering (DLS), and refractive index measurement (RI) were used for additional biophysical characterization (SEC-MALS-SAXS). All samples were centrifuged at 16,000 rpm for 10 min immediately prior to loading on the chromatography system, and were then loaded on a Superdex 200 Increase 10/300 GL column (Cytiva) run by 1260 Infinity II HPLC (Agilent Technologies) at 0.6 ml/min. The flow passed through (in order) the Agilent UV detector, a MALS/DLS detector (DAWN Helios II, Wyatt Technologies), and an RI detector (Optilab T-rEX, Wyatt). The flow then passed through the SAXS flow cell. The flow cell consists of a 1.0 mm ID quartz capillary with ∼20 µm walls. A coflowing buffer sheath is used to separate samples from the capillary walls, helping prevent radiation damage (Kirby et al., 2016). X-ray data were collected using a 150 (h) × 25 (v) (µm) 12 keV beam, and scattering intensity was recorded using an Eiger2 XE 9M (Dectris) photon-counting detector. The detector was placed 3.6 m from the sample giving access to a *q*-range of 0.0029 Å^-1^ to 0.417 Å^-1^, with the momentum transfer vector *q* defined as *q* = (4π⁄λ)sin(θ). Exposures (0.5 s) were acquired every 1 s during elution, and data were reduced using BioXTAS RAW 2.1.1(124). Buffer blanks were created by averaging regions flanking the elution peak and subtracted from exposures selected from the elution peak to create the I(*q*) vs *q* curves used for subsequent analyses. Molecular weights and hydrodynamic radii were calculated from the MALS and DLS data, respectively, using ASTRA 7 software (Wyatt). Further experimental details and data analysis parameters can be found in the Supplementary Tables.

Additional SAXS experiments were performed at the ID7A beamline at the Cornell High Energy Synchrotron Source (CHESS, Ithaca, NY) using similar SEC-MALS-SAXS setup and identical sample preparation procedure. SEC was performed using a Superdex 200 Increase 10/300 GL column (Cytiva) run by a Teledyne Reaxus HPLC pump at 0.6 ml/min. X-ray data were collected using a 250 (h) × 250 (v) (µm) 11.3 keV beam, and scattering intensity was recorded using an Eiger 4M (Dectris) photon-counting detector. The detector was placed 1.78 m from the sample giving access to a *q*-range of 0.0083 Å^-1^ to 0.44 Å^-1^, with the momentum transfer vector *q* defined as *q* = (4π⁄λ)sin(θ). Exposures (1 s) were acquired every 1 s during elution, and data were reduced using BioXTAS RAW 2.2.1(124). Buffer blanks were created by averaging regions preceding the elution peak. Deconvolution by EFA (78) and/or REGALS (79) was performed to separate out the component of interest from other species and accumulated radiation damage. Molecular weights and hydrodynamic radii were calculated from the MALS and DLS data, respectively, using ASTRA 7 software (Wyatt).

### Confocal smFRET analysis

All samples were prepared in buffer (0.16 M NaCl, 20 mM HEPES pH 7.5, 1 mM TCEP, 0.003% Tween 20) and centrifuged at 40,000 rpm for 20 min before the experiment. FRET measurements of freely diffusing single molecules were performed with a confocal microscope [MicroTime 200 (PicoQuant)] as described previously(88, 125). The donor and acceptor were excited by light from 532 nm and 638 nm lasers, respectively. A pulsed interleaved excitation (PIE) setup was used with a pulse rate of 40 MHz to alternate donor and acceptor excitation. PIE reports the status of both donor and acceptor fluorophores by sorting molecules based on relative donor:acceptor stoichiometry (S) and apparent FRET efficiency (E) as described previously (86, 89, 126). Measurements were performed 25-μm deep in the solution using a laser power of ∼15 μW for 30–60 min per sample. Data were recorded using SymPhoTime software 64, version 2.4 (PicoQuant). Data were analyzed with MATLAB-based PAM software (https://pam.readthedocs.io/en/latest/) (127) using a customized profile optimized for our microscope. Signals from single molecules were observed as bursts of fluorescence. Bursts with more than 50 counts were searched with the dual channel burst search (DCBS) algorithm. Integration time was set to 0.5 ms. Appropriate corrections for direct excitation of the acceptor at the donor excitation wavelength (DE), leakage of the donor in the acceptor channel (Lk), and the instrumental factor (g) were determined experimentally using a mixture of dsDNA models with known FRET efficiency and stoichiometry labeled with Cy3 and Cy5: DE=0.05, Lk=0.08, g=0.85.

### Analytical ultracentrifugation (AUC)

PALB2-DBD samples (1 ml) were dialyzed against 0.5 L of buffer (0.16 M NaCl, 25 mM HEPES pH 7.5, 1 mM TCEP) in dialysis bags for 4 hours at room temperature. The sample was centrifuged at 14,000 rpm for 5 min at room temperature and the concentration was measured by Bradford assay. Experiments were performed using a Beckman Coulter Optima AUC. Samples were loaded into cells and centrifuged at 40,000 rpm for 12 h at 21°C. Absorbance was measured at 280 nm for protein samples and at 550 nm for samples with Cy3-labeled DNA every 4 min for a total of 200 scans. Buffer density and viscosity were calculated using the Sednterp program data fitting was performed using the Lamm equation for the continuous c(s) distribution model in Sedfit (128). DNA-only samples were analyzed using a spatial specific volume of 0.55.

## Acknowledgements

We are grateful to Drs. Gregory DeKoster and Carl Frieden for help with CD spectroscopy, Dr. Jaigeeth Deveryshetty for help with AUC, Dr. Sahiti Kupa for help with the preliminary CD spectroscopy experiments, Dr. Samantha Gies for help with EPR spectroscopy, Ian Miller for help with mutagenesis, Laasyapriya Sarva for help with protein purification and activity validation, Dr. Andrea Soranno for helpful discussions, and Dr. Joel Eissenberg for critical reading and help with manuscript preparation. We are also grateful to Dr’s Richard Gillilan, Qingqiu Huang and Steve Meisburger for assistance with SAXS data collection at CHESS. The manuscript was edited by the Scientific Editing Service of the Institute of Clinical and Translational Sciences at Washington University, which is supported by an NIH Clinical and Translational Science Award (UL1 TR002345).

## Funding

This research was supported by Saint Louis University Institute for Drug and Biotherapeutic Innovation and E.A. Doisy Department of Biochemistry and Molecular Biology; Washington University Siteman Cancer Center (SCC), the Foundation for Barnes-Jewish Hospital Siteman Investment Program (SIP) and Institute of Clinical and Translational Sciences JIT Core Usage funding program; and NIH S10 OD030343. SAXS experiments used resources of the Advanced Photon Source, a US Department of Energy (DOE) Office of Science User Facility operated for the DOE Office of Science by Argonne National Laboratory under Contract No. DE-AC02-06CH11357. BioCAT was supported by grant P30 GM138395 from the National Institute of General Medical Sciences of the National Institutes of Health. The content is solely the responsibility of the authors and does not necessarily reflect the official views of the National Institute of General Medical Sciences or the National Institutes of Health. SAXS experiments at CHESS used resources of Center for High-Energy X-ray Sciences (CHEXS), which is supported by the National Science Foundation (BIO, ENG and MPS Directorates) under award DMR-1829070., and the Macromolecular Diffraction at CHESS (MacCHESS) facility, which is supported by award 1-P30-GM124166-01A1 from the National Institute of General Medical Sciences, National Institutes of Health, and by New York State’s Empire State Development Corporation (NYSTAR).

## Author contributions

SK conceptualized the study; YK, MBW, JMR, RD, JH, NP, and SK designed, performed, and analyzed experiments; MBW performed computational modeling of SAXS data; RD designed and analyzed cwEPR; VNU performed sequence analysis; NP helped to design and analyze confocal smFRET; SK wrote the manuscript with input from all authors, supervised the study, and acquired funding.

## Conflict of interest

None declared.

## Data availability

All protein expression plasmids are freely available on request. SAXS scattering data are deposited to SASDBD (https://www.sasbdb.org/) with accession IDs SAS5082 and SAS5083 for high salt and low salt data sets, respectively. The smFRET data in PTU file format (time-correlated single-photon counting data) are available on request.

## Notes

### Competing Interest Statement

The authors have declared no competing interest.

### Summary of Updates

The revised version includes new SAXS data the complex of PALB2-DBD with DNA and revised AUC results.

