## Supplemental Tables an Figures for "The strand exchange domain of tumor suppressor PALB2 is intrinsically disordered and promotes oligomerization-dependent DNA compaction"

**Table S1. SAXS experimental details (APS experiments)**

| **(a) SAXS Sample details** | | |
|---|---|---|
| Sample | PALB2 195W, 500 mM NaCl | PALB2 195W, 160 mM NaCl |
| *MW* from chemical composition (dimer) | 46600 | 46600 |
| Extinction coefficient (280 nm, predicted) | 9970 $M^{-1}$ $cm^{-1}$ (dimer) | 9970 $M^{-1}$ $cm^{-1}$ (dimer) |
| Loading concentration | 110 μM | 100 μM |
| Buffer composition | 20 mM HEPES, 500 mM NaCl, 0.5 mM TCEP pH 7.4 | 20 mM HEPES, 160 mM NaCl, 0.5 mM TCEP pH 7.4 |
| **(b) SAXS data collection parameters** | | |
| Sample | PALB2 195W, 500 mM NaCl | PALB2 195W, 160 mM NaCl |
| Instrument | BioCAT (APS 18ID) with Eiger 2 XE 9M | BioCAT (APS 18ID) with Eiger 2 XE 9M |
| Energy (keV) | 12.0 | 12.0 |
| Beam size ($\mu m^2$) | 150 (h) x 25 (v) | 150 (h) x 25 (v) |
| Camera length (m) | 3.6 | 3.6 |
| $q$-measurement range ($Å^{-1}$) | 0.0029-0.417 | 0.0029-0.417 |
| Absolute Scaling Method | Glassy Carbon, NIST SRM 3600 | Glassy Carbon, NIST SRM 3600 |
| Normalization | Transmitted intensity (beam-stop counter) | Transmitted intensity (beam-stop counter) |
| Configuration | SEC-MALS-SAXS using Superdex 200 Increase 10/300 GL column run by a 1260 Infinity II HPLC (Agilent). UV data was measured in the Agilent, and and MALS-DLS-RI data by DAWN HELEOS-II (17 MALS + 1 DLS channels) and Optilab T-rEX (RI) instruments (Wyatt Technology). SAXS data was measured in a sheath-flow cell (1), effective path length 0.542 mm. | SEC-MALS-SAXS using Superdex 200 Increase 10/300 GL column run by a 1260 Infinity II HPLC (Agilent). UV data was measured in the Agilent, and and MALS-DLS-RI data by DAWN HELEOS-II (17 MALS + 1 DLS channels) and Optilab T-rEX (RI) instruments (Wyatt Technology). SAXS data was measured in a sheath-flow cell (1), effective path length 0.542 mm. |
| Experimental temperature (° C) | 23 | 23 |

**Table S2: SAXS data processing and analysis (APS experiments)**

| **(a) SAXS data processing and structural parameters** | | |
|---|---|---|
| Sample | PALB2 195W, 500 mM NaCl | PALB2 195W, 160 mM NaCl |
| Buffer range (frames) | 50-250 | 544-674 |
| Sample range (frames) | 1271-1296 | 1414-1464 |
| **Guinier Analysis** | | |
| *I(0)* | 0.0234 ± 0.000046 | 0.00770 ± 0.000023 |
| $R_g$ (Å) | 54.88 ± 0.22 | 47.48 ± 0.24 |
| $qR_g$ range | 0.241-1.142 | 0.176-1.290 |
| $R^2$ (fit) | 0.992 | 0.982 |
| ***P*I analysis (GNOM)** | | |
| *I(0)* | 0.0234 ± 0.000070 | 0.00781 ± 0.000033 |
| $R_g$ (Å) | 58.35 ± 0.44 | 50.98 ± 0.45 |
| $d_{max}$ (Å) | 254 | 213 |
| $q$-range (Å$^{-1}$) | 0.0044-0.417 | 0.0038-0.417 |
| $\chi^2$ /total estimate | 0.852/0.654 | 0.857/0.0.683 |
| **MFF analysis** | | |
| $R_g$ (Å) | 55.02 ± 0.07 | 49.10 ± 0.12 |
| $\nu$ | 0.520 ± 0.01 | 0.472 ± 0.002 |
| $\chi_r^2$ | 0.993 | 0.839 |
| *MW*, kDa ($V_C$) | 67.7 | 69.5 |
| **(b) Software employed for SAS reduction and analysis** | | |
| SAXS data reduction | Radial averaging; frame comparison, averaging, and subtraction done using BioXTAS RAW 2.1.1 (2) | Radial averaging; frame comparison, averaging, and subtraction done using BioXTAS RAW 2.1.1 (2) |
| Basic analysis: Guinier, MW, P(r) | Guinier fit and M.W. using BioXTAS RAW, P(r) function using GNOM (3). RAW uses MoW and Vc M.W. methods (4,5) | Guinier fit and M.W. using BioXTAS RAW, P(r) function using GNOM (3). RAW uses MoW and Vc M.W. methods (4,5) |
| MALS-DLS-RI analysis | Astra 7 (Wyatt) | Astra 7 (Wyatt) |

**Table S3. SAXS experimental details (CHESS experiments)**

| **(a) SAXS Sample details** | | |
|---|---|---|
| Sample | PALB2 L24A, 160 mM NaCl | PALB2 +dT50, 160 mM NaCl |
| *MW* from chemical composition | 23300 | 61700 |
| Extinction coefficient (280 nm, predicted) | 2980 $M^{-1}$ $cm^{-1}$ (monomer) | 9970 $M^{-1}$ $cm^{-1}$ (protein)/ 405600 $M^{-1}$ $cm^{-1}$ (DNA) |
| Loading concentration | 220 μM | 160 μM (protein); 2:1 protein:DNA |
| Buffer composition | 20 mM HEPES, 160 mM NaCl, 0.5 mM TCEP pH 7.5 | 20 mM HEPES, 160 mM NaCl, 0.5 mM TCEP pH 7.5 |

| **(b) SAXS data collection parameters** | | |
|---|---|---|
| Sample | PALB2 L24A, 160 mM NaCl | PALB2 +dT50, 160 mM NaCl |
| Instrument | CHESS ID7A with Eiger 4M | CHESS ID7A with Eiger 4M |
| Energy (keV) | 11.3 | 11.3 |
| Beam size ($\mu m^2$) | 250 (h) x 250 (v) | 250 (h) x 250 (v) |
| Camera length (m) | 1.78 | 1.78 |
| *q*-measurement range ($Å^{-1}$) | 0.0083-0.44 | 0.0083-0.44 |
| Normalization | Transmitted intensity (beam-stop counter) | Transmitted intensity (beam-stop counter) |
| Configuration | SEC-MALS-SAXS using Superdex 200 Increase 10/300 GL column run by a Teledyne Reaxus HPLC pump. UV data was measured with an ATKA Pure MWD detector, and and MALS-DLS-RI data by DAWN HELEOS-II (17 MALS + 1 DLS channels) and Optilab T-rEX (RI) instruments (Wyatt Technology). | SEC-MALS-SAXS using Superdex 200 Increase 10/300 GL column run by a Teledyne Reaxus HPLC pump. UV data was measured with an ATKA Pure MWD detector, and and MALS-DLS-RI data by DAWN HELEOS-II (17 MALS + 1 DLS channels) and Optilab T-rEX (RI) instruments (Wyatt Technology). |
| Experimental temperature (° C) | 23 | 23 |

**Table S4: SAXS data processing and analysis (CHESS experiments)**

| **(a) SAXS data processing and structural parameters** | | |
|---|---|---|
| Sample | PALB2 L24A, 160 mM NaCl | PALB2 +dT50, 160 mM NaCl |
| Buffer range (frames) | 402-564 | 186-325 |
| Sample range (frames) | 1300-1583 (REGALS) | 1154-1431 (EFA) |
| **Guinier Analysis** | | |
| *I(0)* | 5.09 ± 0.0504 | 4.26 ± 0.023 |
| $R_g$ (Å) | 52.30 ± 0.70 | 46.11 ± 0.36 |
| $qR_g$ range | 0.688-1.268 | 0.384-1.296 |
| $R^2$ (fit) | 0.946 | 0.966 |
| ***P*I analysis (GNOM)** | | |
| *I(0)* | 5.136 ± 0.0474 | 4.278 ± 0.0191 |
| $R_g$ (Å) | 56.47 ± 0.78 | 47.88 ± 0.32 |
| $d_{max}$ (Å) | 216 | 176 |
| *q*-range ($Å^{-1}$) | 0.0132-0.443 | 0.0083-0.443 |
| $\chi^2$ /total estimate | 1.102/0.593 | 0.885/0.715 |
| **MFF analysis** | | |
| $R_g$ (Å) | 48.81 ± 0.21 | 47.21 ± 0.19 |
| $\nu$ | 0.541 ± 0.003 | 0.384 ± 0.002 |
| $\chi_r^2$ | 1.950 | 0.959 |
| *MW*, kDa ($V_C$) | 49.6 | 110.8 |
| **(b) Software employed for SAS reduction and analysis** | | |
| SAXS data reduction | Radial averaging; frame comparison, averaging, and subtraction done using BioXTAS RAW 2.1.1 (Hopkins *et al.* 2017) | Radial averaging; frame comparison, averaging, and subtraction done using BioXTAS RAW 2.1.1 (Hopkins *et al.* 2017) |
| Basic analysis: Guinier, MW, P(r) | Guinier fit and M.W. using BioXTAS RAW, P(r) function using GNOM (3). RAW uses MoW and Vc M.W. methods (4,5). | Guinier fit and M.W. using BioXTAS RAW, P(r) function using GNOM (3). RAW uses MoW and Vc M.W. methods (4,5). |
| MALS-DLS-RI analysis | Astra 7 (Wyatt) | Astra 7 (Wyatt) |

**Table S5: Ensemble Fit Statistics**

| **(a) SASSIE Ensemble (unguided)** | ***500 mM NaCl*** | ***160 mM NaCl*** |
|---|---|---|
| Structures in starting pool | 34,919 | 34,919 |
| Fit $\chi^2$ value | 1.006 | 2.034 |
| $R_{flex}$ (random) / $R_{sigma}$ | 64.36% (~ 80.11%) / 1.06 | 56.40% (~ 80.11%) / 1.37 |
| Final ensemble $R_g$ (Å) | 56.67 | 58.19 |
| Final ensemble $D_{max}$ (Å) | 210.38 | 198.26 |
| Structures in final ensemble | 6 | 7 |
| **(b) SASSIE Ensemble (*Rg* guided)** | ***500 mM NaCl*** | ***160 mM NaCl*** |
| Structures in starting pool | N/A | 34,819 |
| Fit $\chi^2$ value | N/A | 1.320 |
| $R_{flex}$ (random) / $R_{sigma}$ | N/A | 62.68% (~ 66.13%) / 1.35 |
| Final ensemble $R_g$ (Å) | N/A | 50.84 |
| Final ensemble $D_{max}$ (Å) | N/A | 185.04 |
| Structures in final ensemble | N/A | 5 |
| **(c) SASSIE Ensemble (unguided, monomer)** | | |
| Structures in starting pool | N/A | 38,042 |
| Fit χ2 value | N/A | 1.395 |
| $R_{flex}$ (random) / $R_{sigma}$ | N/A | 81.39% (~ 80.39%) / 1.46 |
| Final ensemble $R_g$ (Å) | N/A | 58.23 |
| Final ensemble $D_{max}$ (Å) | N/A | 195.65 |
| Structures in final ensemble | N/A | 6 |

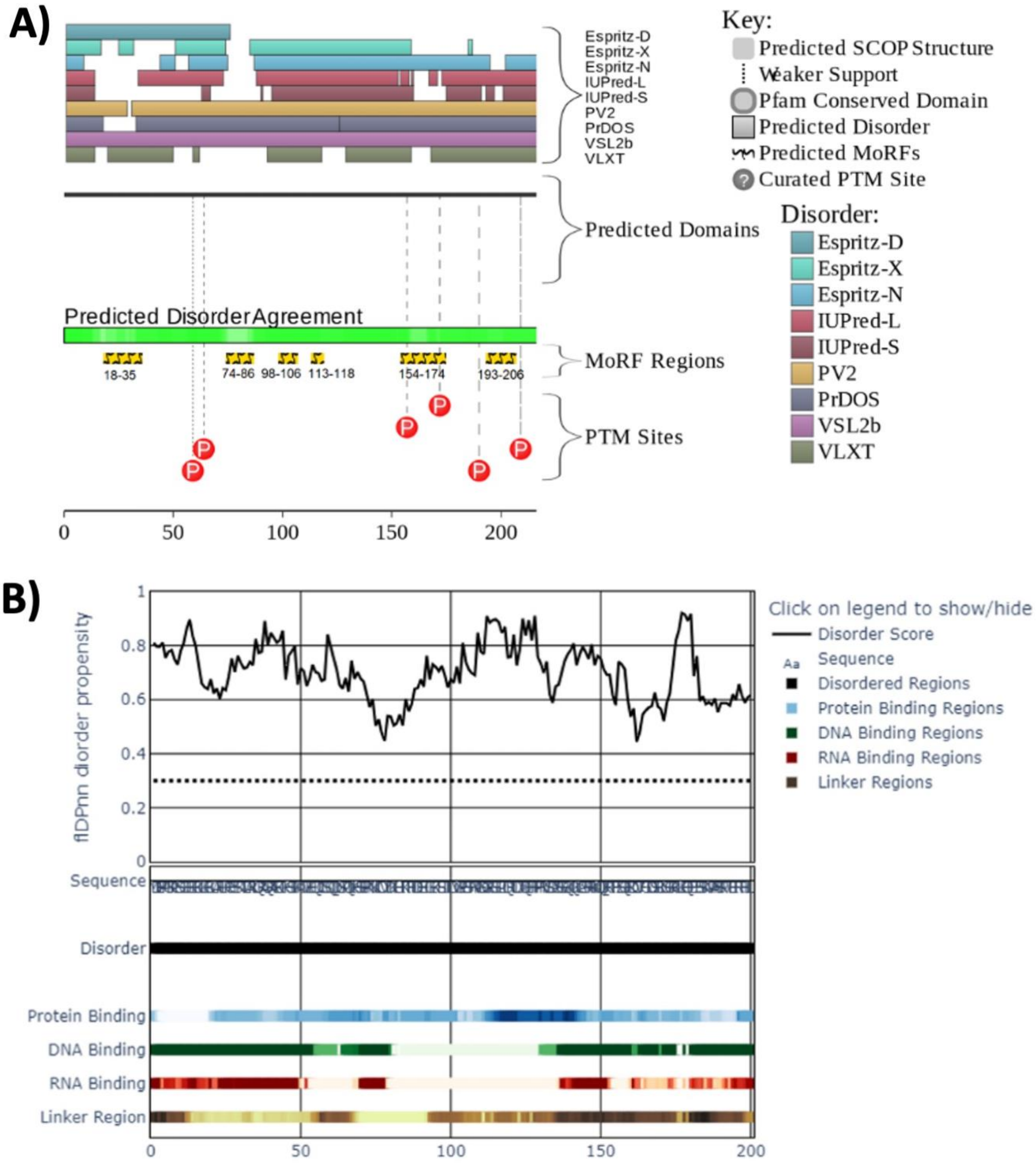


**Supplementary Figure S1**. **PALB2-DBD amino acid sequence analysis. A)** Functional disorder profile generated by the $D^2P^2$ platform (https://d2p2.pro/); **B)** Functional disorder profile generated by the flDPnn platform (http://biomine.cs.vcu.edu/servers/flDPnn/)

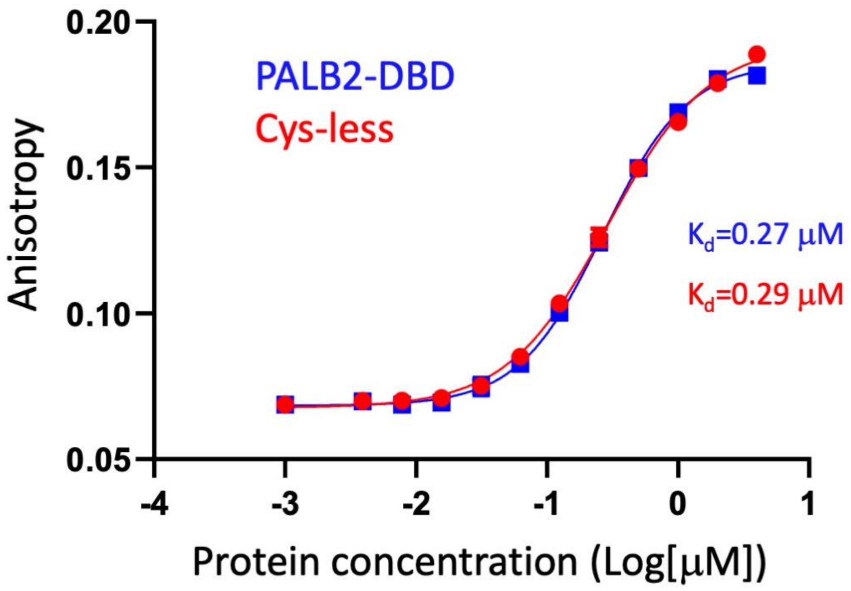


**Supplementary Figure S2**. Binding of FAM-dT50 by PALB2-DBD (blue) and with mutant with all cysteins mutated to alanines (red). The measurements were performed exactly as described in (32) with final concentration of FAM-dT50 of 10 nM in 0.16 M NaCl, 25 mM HEPES 7.5, 1 mM TCEP at 20°C in a plate reader. Each point is an average of three measurements.

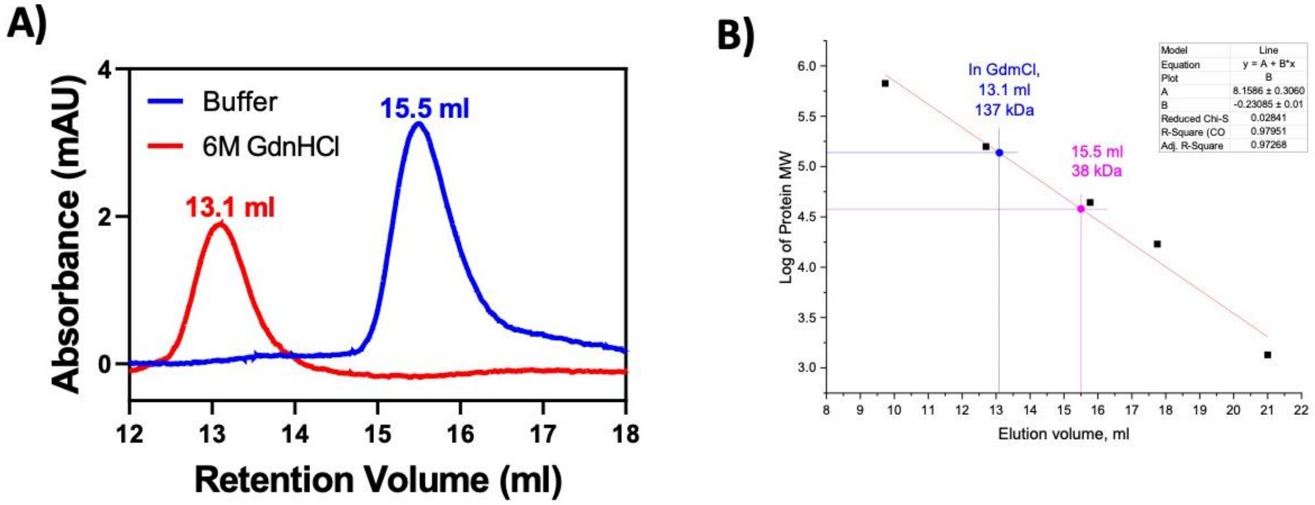


**Supplementary Figure S3**. Size exclusion chromatography of Δ40-PALB2. **A)** Elution profile in the DNA-binding buffer is shown in blue and that in GndHCl in red. Volumes of peaks elution are shown on top. **B)** Estimation of an apparent molecular weight for each peak based on elution volumes of protein markers (BioRad, cat #1511901).

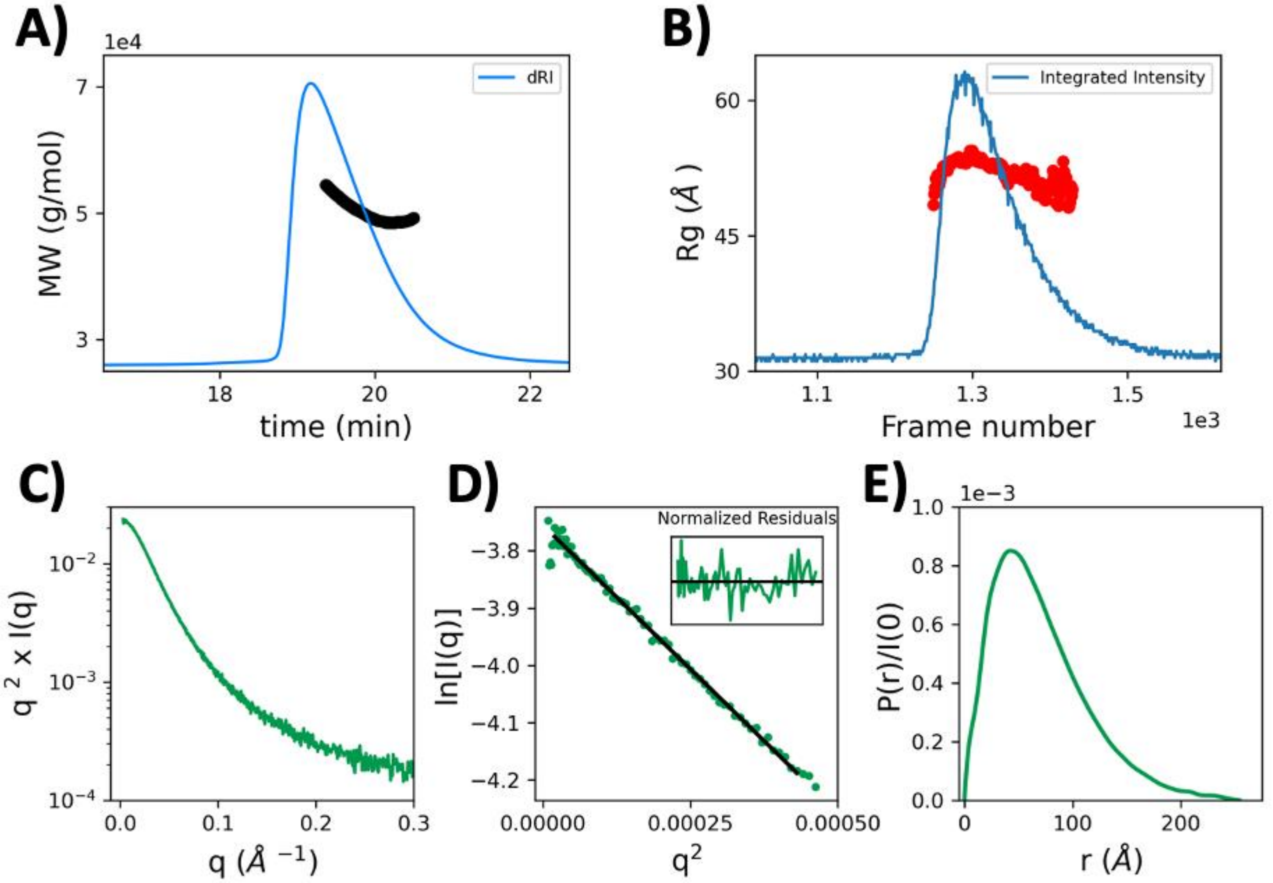


**Supplementary Figure S4: SEC-MALS SAXS of PALB2-DBD in 500 mM NaCl buffer. (A)** Calculated MALS molecular weight across the elution peak (as measured by differential refractive index, dRI). Calculated MW's for the main part of the peak were as expected for a monomer, ~45-50 kDa. Calculation at the tails of the peak unreliable. **(B)** SAXS *Rg* values plotted across the elution peak (as displayed by integrated intensity). The sample appeared largely monodisperse, with potential weak concentration effects at the tails of the peaks. **(C)** SAXS profile, shown as a semilog plot. **(D)** Guinier fit for the data shown in **C**. No trends in residuals are observed across the fit region. **(E)** IFT (calculated in GNOM (3)) for the dataset shown in **A**. The shape is indicative of a somewhat extended conformation.

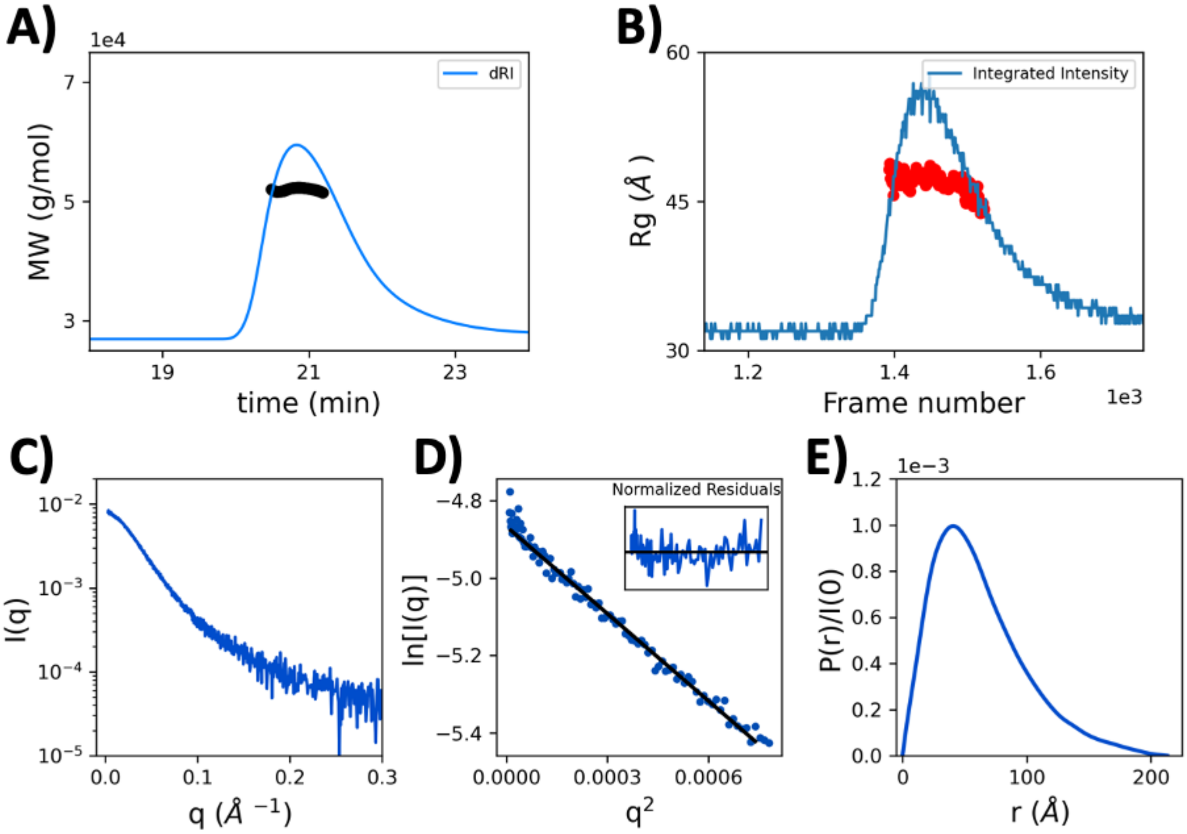


**Supplementary Figure S5: SEC-MALS SAXS of PALB2-DBD in 160 mM NaCl buffer. (A)** Calculated MALS molecular weight across the elution peak (as measured by differential refractive index, dRI). Calculated MW's for the main part of the peak were close to as expected for a monomer, ~52 kDa. Calculation at the tails of the peak unreliable. **(B)** SAXS *Rg* values plotted across the elution peak (as displayed by integrated intensity). The sample appeared largely monodisperse, with potential weak concentration effects at the tails of the peaks. **(C)** SAXS profile, shown as a semilog plot. **(D)** Guinier fit of the data shown in **C**. No significant trends in residuals are observed across the fit region. **(E)** IFT (calculated in GNOM (3)) for the dataset shown in B. The shape is indicative of a more compact conformation than the high salt condition.

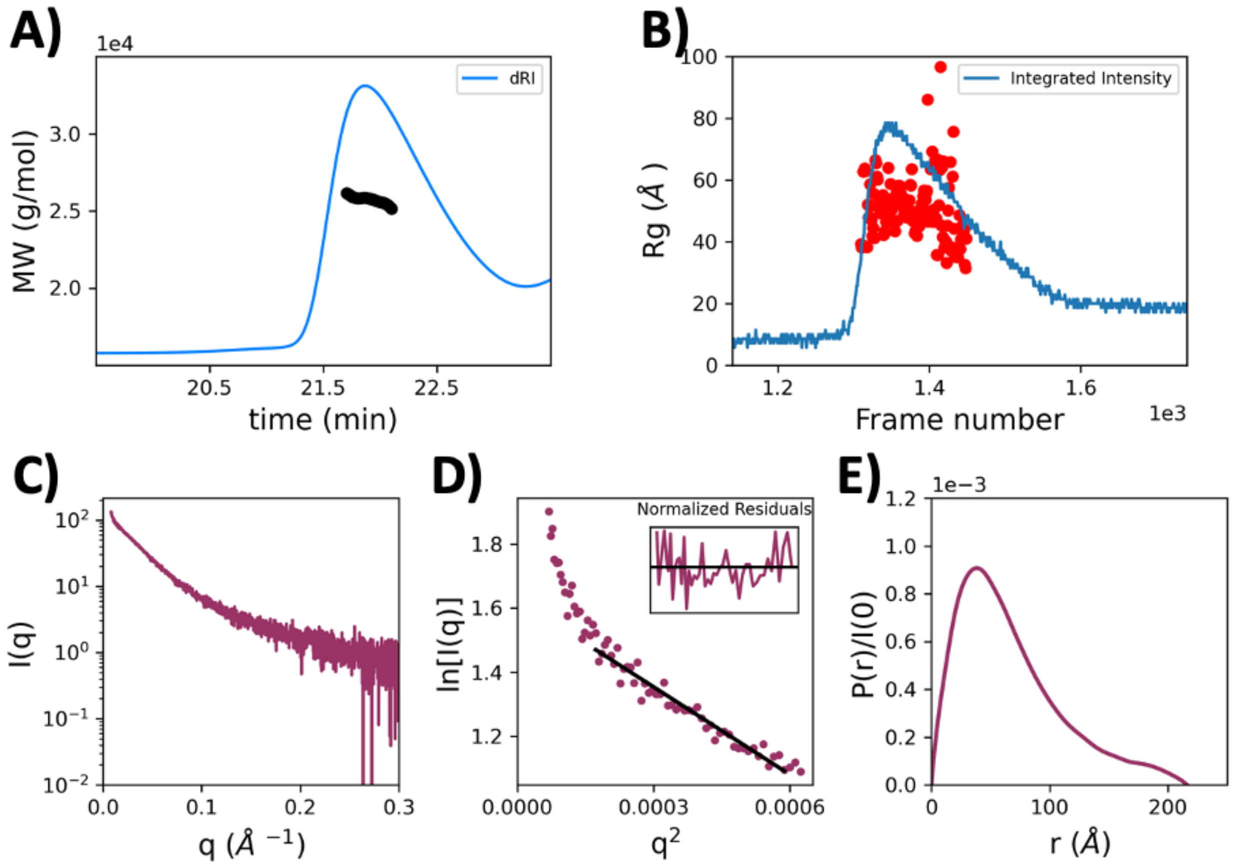


**Supplementary Figure S6: SEC-MALS SAXS of PABL2 L24A (A)** Calculated MALS molecular weight across the elution peak (as measured by differential refractive index, dRI). Calculated MW's for the main part of the peak were similar to expected for a monomer, ~26 kDa. Calculation at the tails of the peak unreliable. **(B)** SAXS *Rg* values plotted across the elution peak (as displayed by integrated intensity). The sample appeared relatively monodisperse, with some accumulated radiation damage visible. **(C)** SAXS profile, shown as a semilog plot. **(D)** Guinier fit for the data shown in **C**. Some radiation damage is visible. **(E)** IFT (calculated in GNOM (3)) for the dataset shown in A. The shape is indicative of an extended conformation.

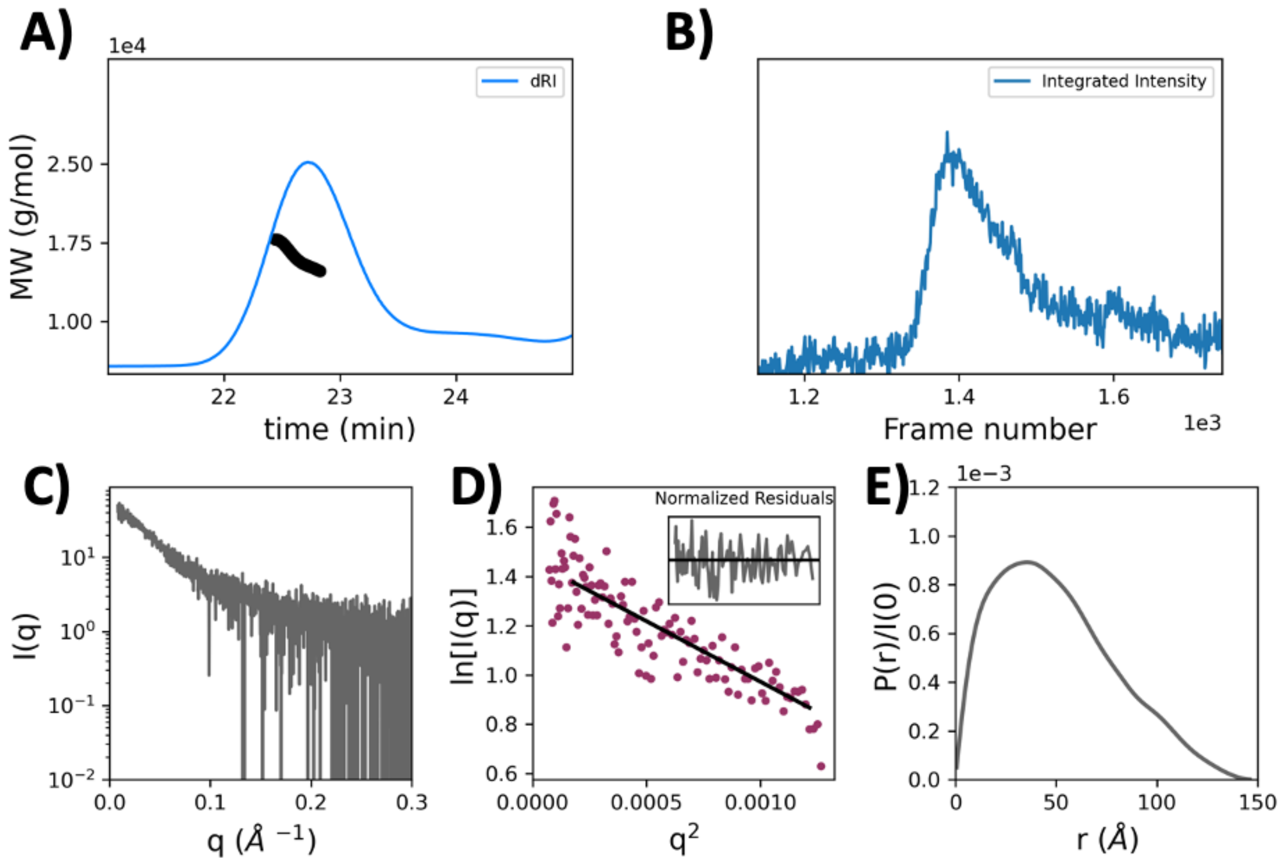


**Supplementary Figure S7: SEC-MALS SAXS of dT50 (A)** Calculated MALS molecular weight across the elution peak (as measured by differential refractive index, dRI). Calculated MW's for the main part of the peak were similar to expected for a monomer, ~15-17 kDa. Calculation at the tails of the peak unreliable. **(B)** SAXS integrated intensity; *Rg* calculation for individual profiles was unreliable due to the low signal in the dataset. The sample appeared largely monodisperse, with potential weak concentration effects at the tails of the peaks. **(C)** SAXS profile, shown as a semilog plot. **(D)** Guinier fit for the data shown in **C**. No trends in residuals are observed across the fit region. **(E)** IFT (calculated in GNOM (3)) for the dataset shown in **B**. The shape is indicative of a rod-shaped structure.

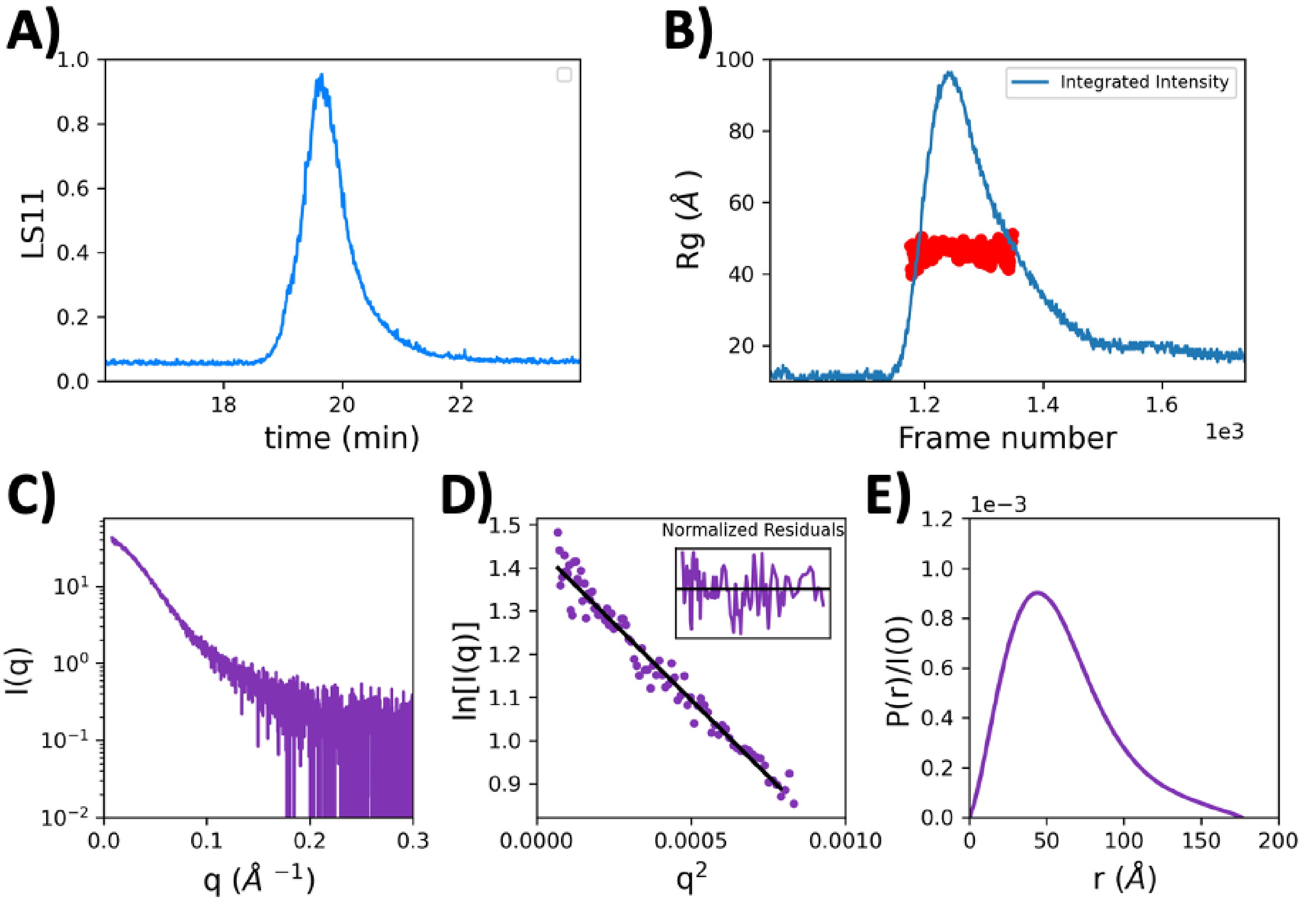


**Supplementary Figure S8: SEC-MALS SAXS of PABL2-dT50 complex (A)** Light scattering channel 11 across the elution peak, from the MALS data collection (the dRI signal was saturated). MW calculations proved unreliable. **(B)** SAXS *Rg* values plotted across the elution peak (as displayed by integrated intensity). **(C)** SAXS profile, shown as a semilog plot. **(D)** Guinier fit for the data shown in **C**. No trends in residuals are observed across the fit region. **(E)** IFT (calculated in GNOM (3)) for the dataset shown in **B**. The shape is indicative of a fairly compact, globular protein potentially with some extended regions.

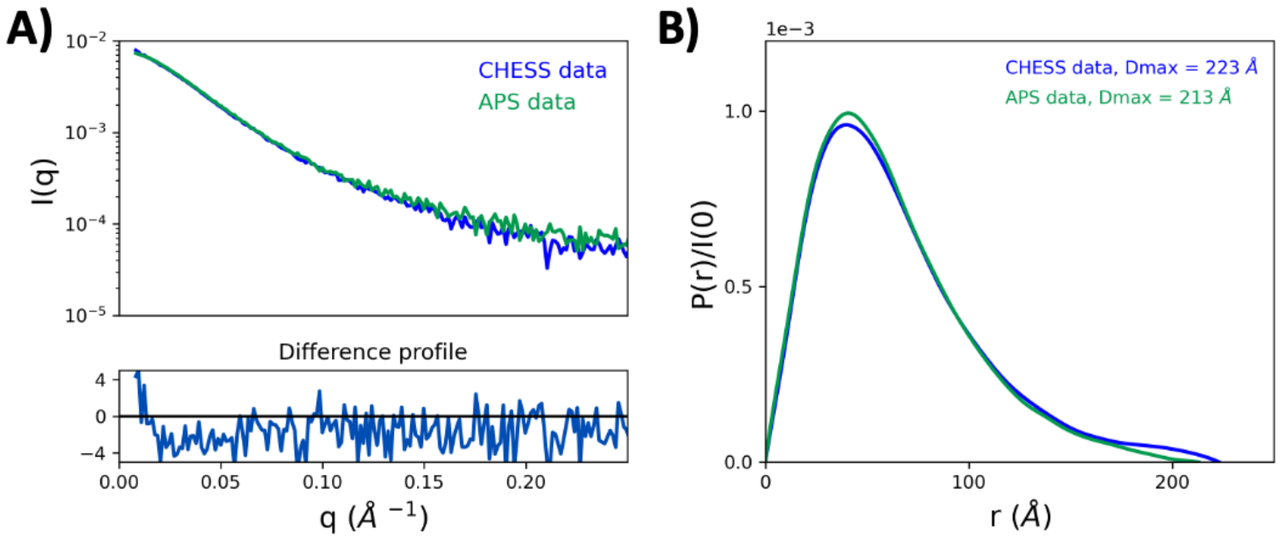


**Supplementary Figure S9: Comparison of PABL2 SAXS datasets (A)** *I(0)*-normalized datasets for PALB2 in 160 mM NaCl, collected at CHESS ID7A (blue) and APS 18ID (green). The datasets overlay well, and appear to represent the same species. The difference profile (bottom) suggests the only meaningful difference is at low-*q*, likely due to increased radiation damage in the CHESS dataset. **(B)** IFT's (calculated in GNOM (3)) for the datasets shown in **A**.

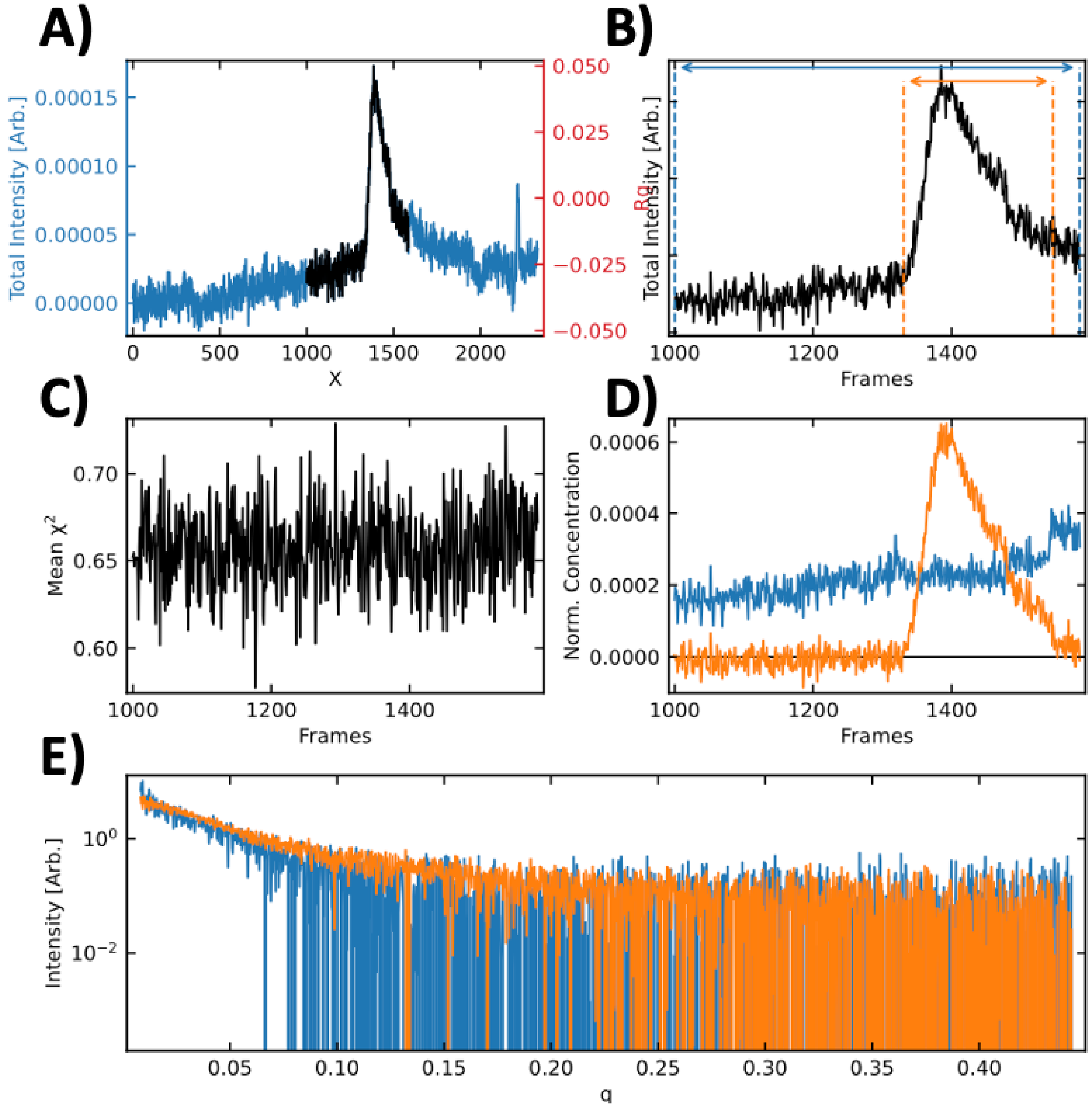


**Supplementary Figure S10: REGALS (6) deconvolution of dT50 SAXS data.** The dataset was separated into a main component (dT50, orange) and an accumulating radiation damage component (blue). **(A)** the deconvoluted region (black) overlayed with the complete dataset (blue). **(B)** peak ranges corresponding to the two components. **(C)** Chi2 value plotted over the deconvoluted region. **(D)** Deconvoluted elution profiles and **(E)** corresponding SAXS profiles for the two components. Analysis was performed only on the main component (orange).

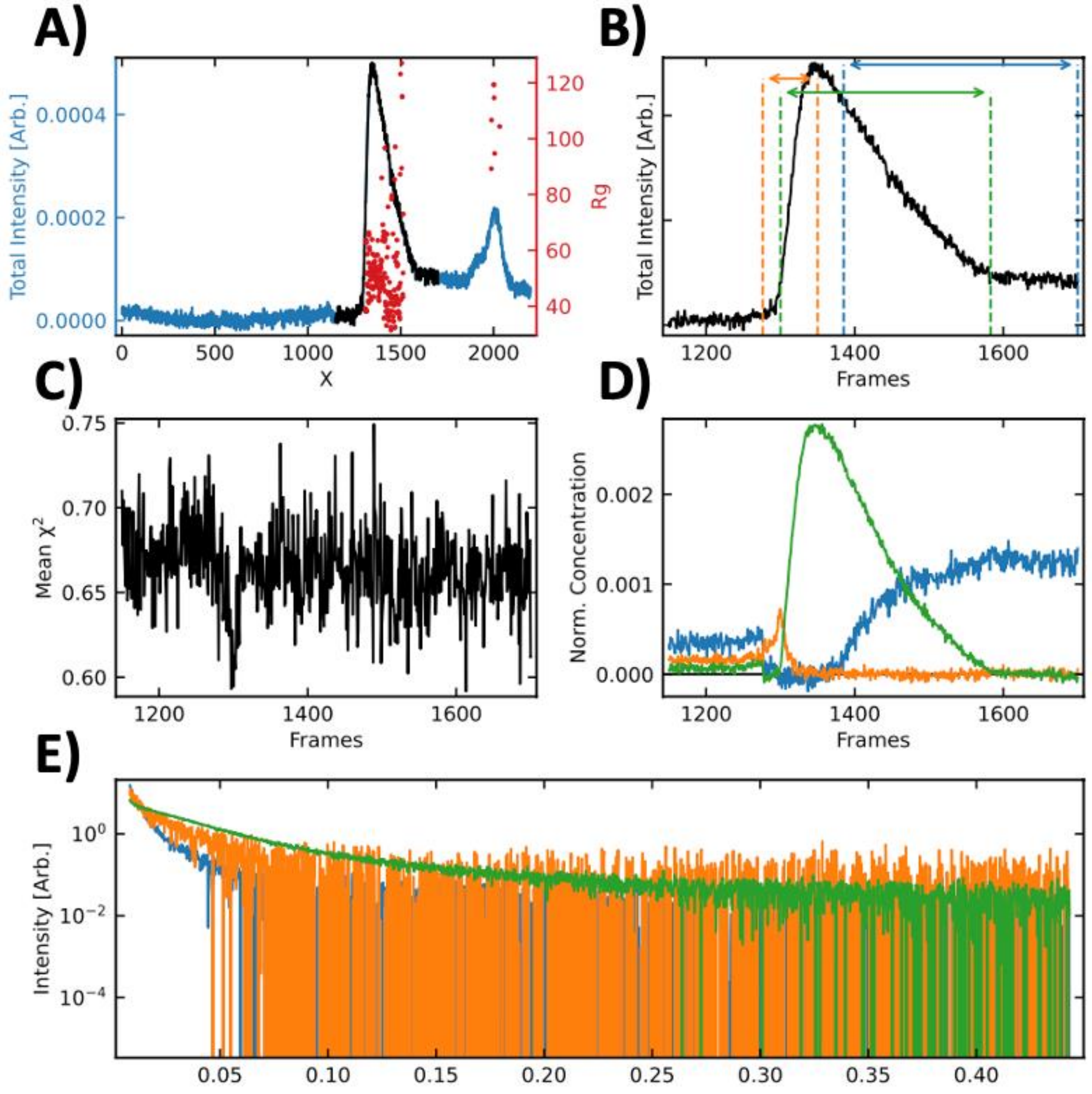


**Supplementary Figure S11: REGALS (6) deconvolution of L24A SAXS data.** The dataset was separated into a main component (L24A monomer, green), an aggregate or large contaminant component (orange) and an accumulating radiation damage component (blue). **(A)** the deconvoluted region (black) overlayed with the complete dataset (blue). **(B)** peak ranges corresponding to the two components. **(C)** Chi2 value plotted over the deconvoluted region. **(D)** Deconvoluted elution profiles and **(E)** corresponding SAXS profiles for the three components. Analysis was performed only on the main component (green).

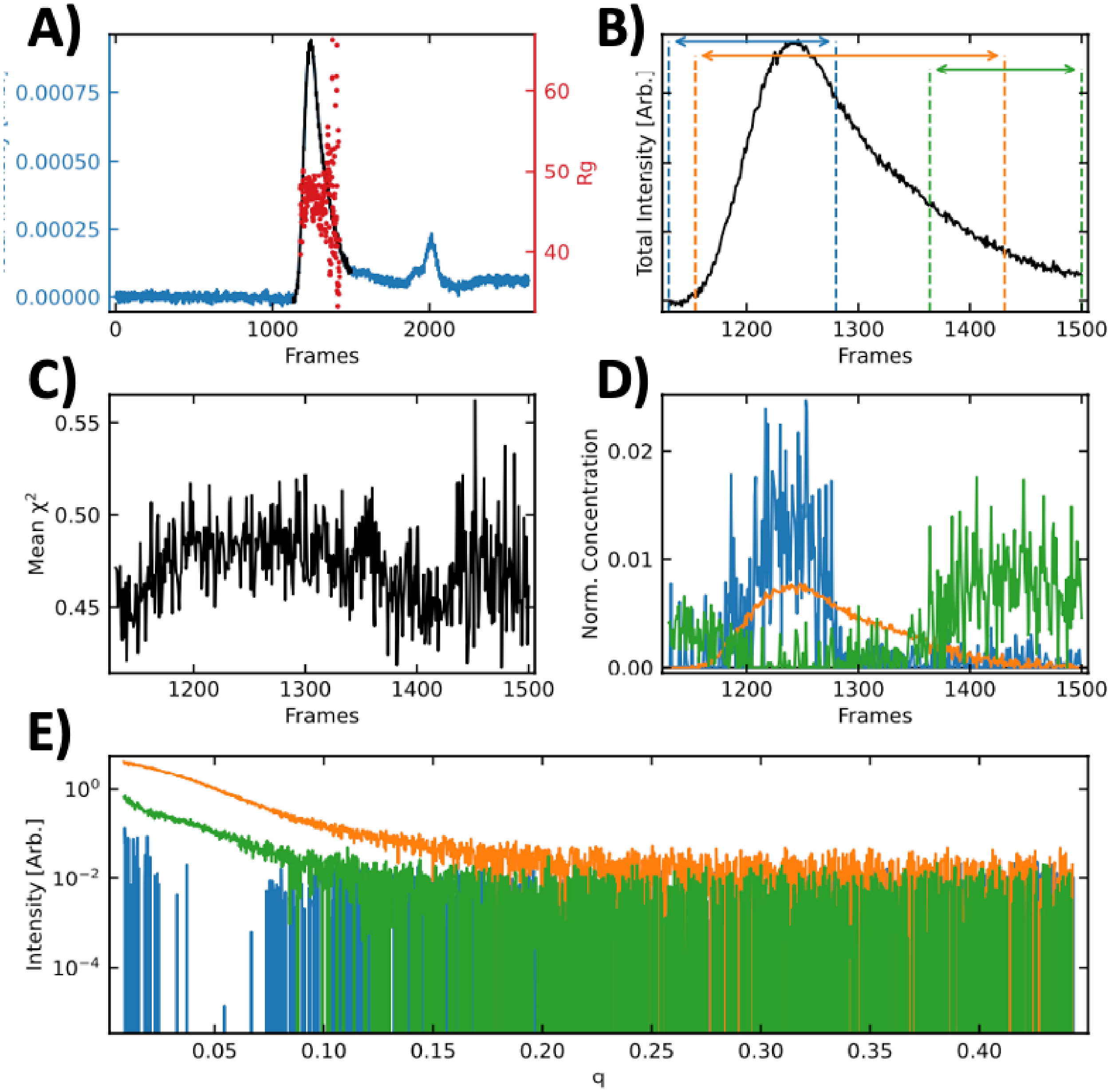


**Supplementary Figure S12: EFA (7) deconvolution of PABLS2-dT50 complex SAXS data.** The dataset was separated into a main component (protein-DNA complex, green), an aggregate or large contaminant component (blue) and a smaller component (blue), potentially protein without DNA. **(A)** the deconvoluted region (black) overlayed with the complete dataset (blue). **(B)** peak ranges corresponding to the two components. (C) Chi2 value plotted over the deconvoluted region. **(D)** Deconvoluted elution profiles and **(E)** corresponding SAXS profiles for the three components. Analysis was performed only on the main component (orange).

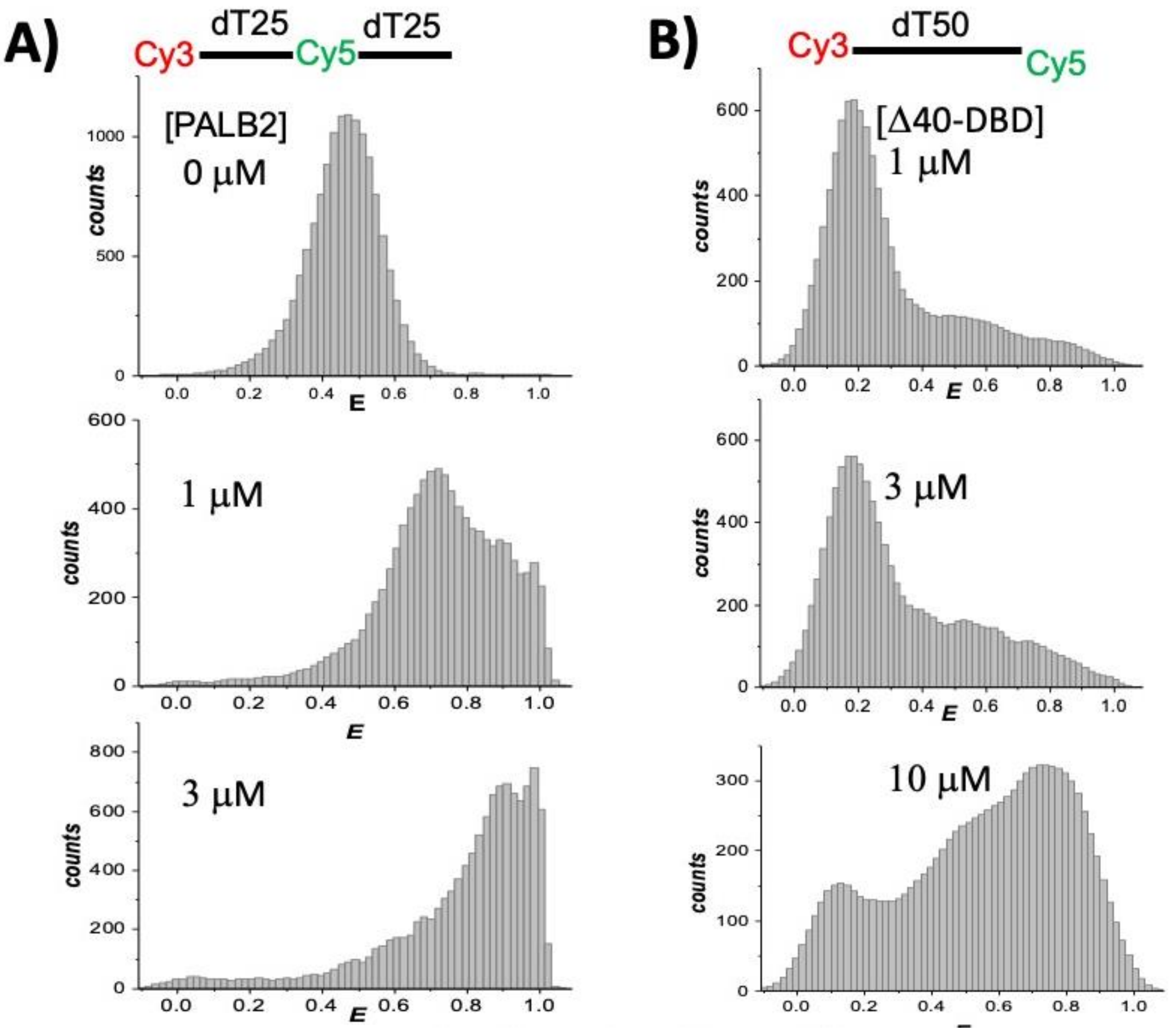


**Supplementary Figure S13. Single-molecule FRET measurements of ssDNA compaction. (A)** FRET histograms of 100 pM $dT_{50}$ with Cy3 placed at the 5′ end and Cy5 attached at position 25 of the $dT_{50}$ substrate alone and in the presence of 1 μM and 3 μM PALB2-DBD. **(B)** FRET histograms of Cy3-$dT_{50}$-Cy5 titrated by Δ40-DBD. Protein concentrations are shown on each graph.

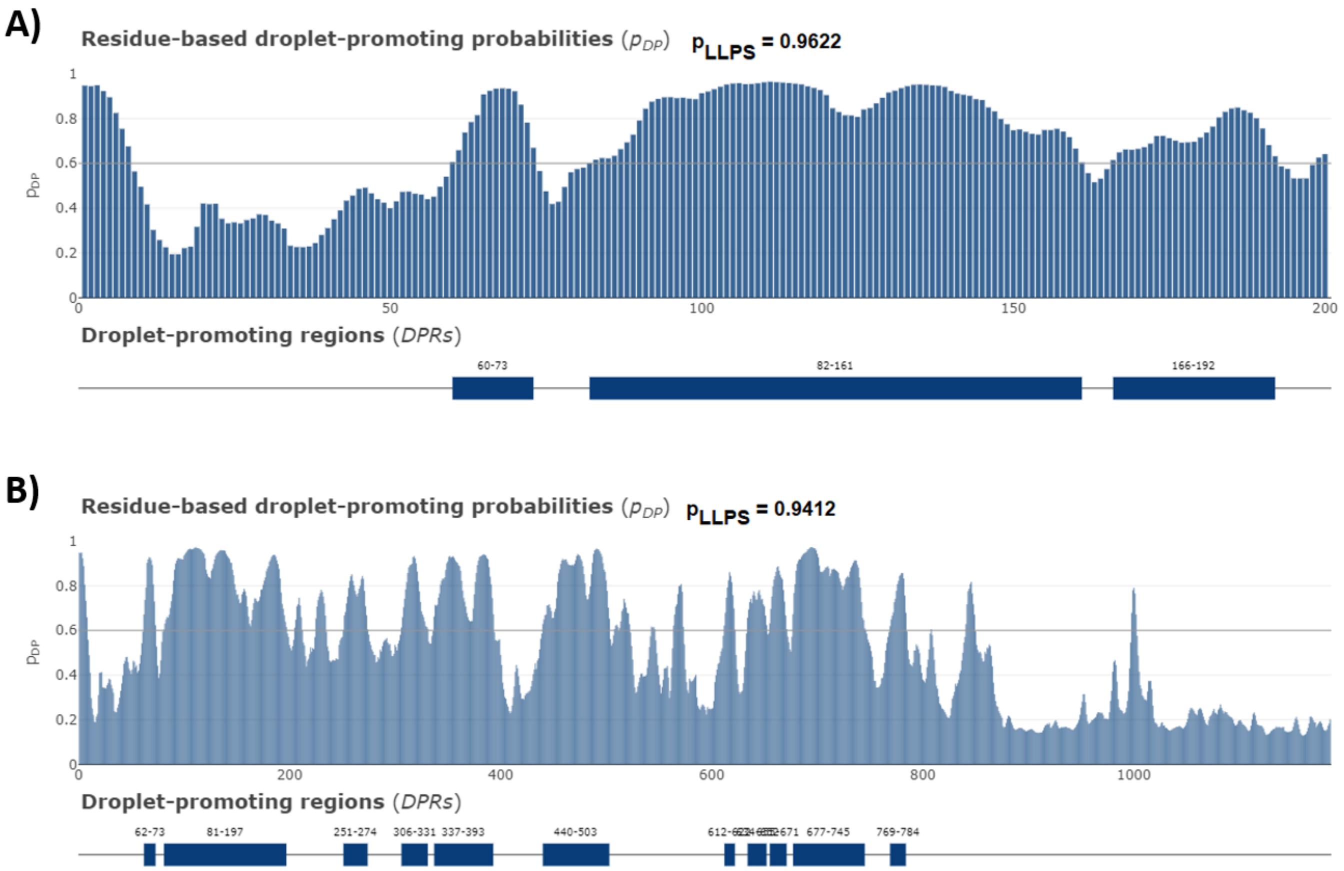


**Supplementary Figure S14**. Distributions of the per-residue droplet-promoting probabilities ($p_{DP}$) within the amino acid sequences of PALB2-DBD (**A)** and PALB2 (**B)** as evaluated by FuzDrop platform (https://fuzdrop.bio.unipd.it/predictor). Positions of the droplet-promoting regions are shown as blue boxes at the bottom of the corresponding plots.

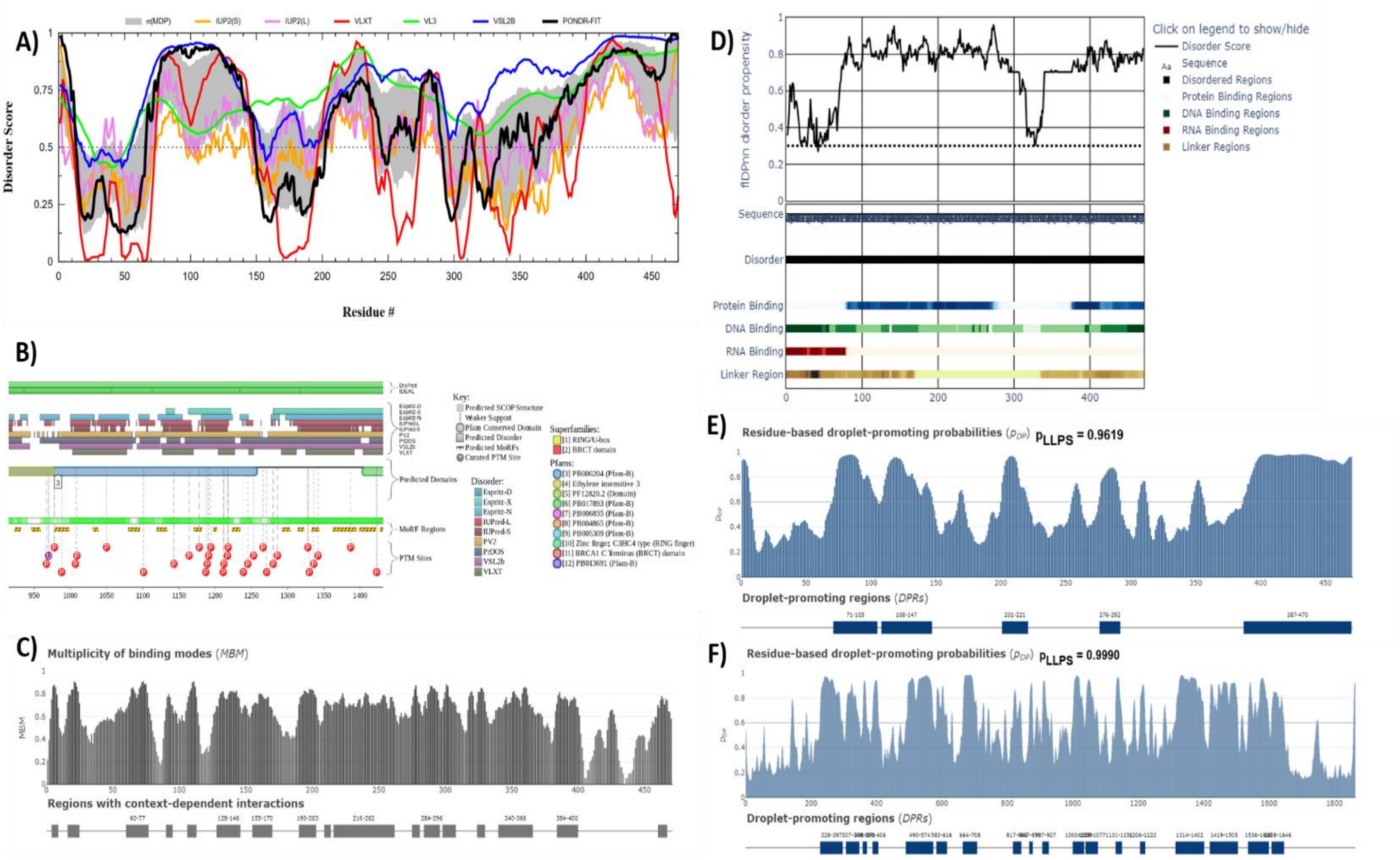


**Supplementary Figure S15**. Intrinsic disorder-centric analysis of the BRCA1-DBD (residues 930-1400). **A)** Disorder profile generated by RIDAO; **B)** Functional disorder profile generated by $D^2P^2$; **C)** Multiplicity of binding modes evaluated by FuzDrop;  **D)** Functional disorder profile generated by the flDPnn platform; **E)** Distributions of the per-residue droplet-promoting probabilities ($p_{DP}$); **F)** Distributions of the per-residue droplet-promoting probabilities ($p_{DP}$) within the full-length human BRCA1.
